# The Causal Pivot: A Structural Approach to Genetic Heterogeneity and Variant Discovery in Complex Diseases

**DOI:** 10.1101/2024.11.03.621751

**Authors:** Chad Shaw, Christopher Williams, Tao Tao Tan, Nicholas Di, Joshua Shulman, John W. Belmont

## Abstract

We present the Causal Pivot (CP) as a structural causal model (SCM) for analyzing genetic heterogeneity in complex diseases. The CP leverages one established causal factor to detect the contribution of a second suspected cause. Specifically, polygenic risk scores (PRS) serve as known causes, while rare variants (RV) or RV ensembles are evaluated as candidate causes. The CP incorporates outcome-induced association by conditioning on disease status. We derive a conditional maximum likelihood procedure for binary and quantitative traits and develop the Causal Pivot Likelihood Ratio Test (CP-LRT) to detect causal signals.

Through simulations, we demonstrate the CP-LRT’s robust power and superior error control compared to alternatives. We apply the CP-LRT to UK Biobank (UKB) data, analyzing three exemplar diseases: hypercholesterolemia (HC, LDL-c ≥ 4.9 mmol/L; n_c_=24,656), breast cancer (BC, ICD10 C50; n_c_=12,479), and Parkinson’s disease (PD, ICD10 G20; n_c_=2,940). For PRS, we utilize UKB-derived values, and for RVs, we analyze ClinVar pathogenic/likely pathogenic variants and loss-of-function mutations in disease-relevant genes: *LDLR* for HC, *BRCA1* for BC, and *GBA* for PD. Significant CP-LRT signals were detected for all three diseases. Cross-disease and synonymous variant analyses serve as controls. We further develop ancestry adjustment using matching and inverse probability weighting, and we extend the CP to examine oligogenic burden in the lysosomal storage pathway for PD. The CP reveals an approach to address heterogeneity and is an extensible method for inference and discovery in complex disease genetics.

## Introduction

Causal heterogeneity, where root causes of disease vary among affected individuals, is a signature characteristic of complex disease. Affected individuals may reach the disease phenotype through distinct etiologies which may be influenced by alternative genetic factors. Although the causes of disease vary, they ultimately lead to shared pathophysiology or disease state. Moreover, complex disease genetic architectures are known to vary from monogenic to highly polygenic.^1,2^ Oligogenic inheritance, environmental phenocopies, and gene X environment interactions are also thought to contribute to complex disease etiologies. This causal heterogeneity impacts both prevention and management because distinct causes may benefit from different approaches for effective intervention or risk mitigation.^3^

Genome wide association studies (GWAS) in complex disease are well developed for both rare and common DNA variants^4,5^, but integration across the allele frequency spectrum remains challenging. Moreover, non-familial association approaches are typically case-control designs and may not be applicable to all settings.^6,7^ Although GWAS has tools for both rare and common variants, almost all studies are marginal analyses that treat each genetic locus as a separate testing unit; results are summarized with Manhattan plots. The marginal approach avoids confronting heterogeneity with respect to alternative genetic factors differentially present or absent in individuals. Consequently, these methods do not classify individuals into causal groups or subtypes. Resolution of heterogeneity is desirable to avoid overgeneralizing results across different subpopulations and to identify which specific causal mechanism or pathway is potentially responsible in individual cases.^8^

Genetic variants -- while exhibiting allelic heterogeneity, pleiotropy and potential confounding with ancestry and environment -- are explicitly considered to be causal factors that drive downstream disease relevant processes. Consequently, methods developed for causalinference and structural causal modeling (SCM) in fields such as epidemiology and econometrics ^9,10^ have been adapted to problems in genetics. Perhaps the most successful example is Mendelian randomization (MR)^11–14^, which uses genetic variants as instrumental variables to evaluate potential causal biomarkers and endophenotypes for common diseases. We reasoned causal analysis methods could have broader applications in genetics^15–19^ and could offer a potential approach to reframe inquiry into heterogeneity in complex disease genetics.

We hypothesized that SCM could serve as a framework to consider multiple genetic factors differentially present or absent in individuals in order to examine the underlying mechanistic heterogeneity of complex disease. We adopt the strategy to model common variant and rare genetic contributions to disease as separate causal contributions where disease outcome is their common effect. We encapsulate the common variant contribution to disease in a polygenic risk score (PRS). The graphical structure presents a collider pattern between PRS and RV. The general concept that low PRS can be used to prioritize affected individuals for RV sequencing or to classify them as having monogenic disorders has been noted^20,21^. More generally, the use of colliders or v-structures to infer causal graphs is well known.^22–24^ However, more detailed statistical analysis is still needed to elevate and generalize the procedure, and recognition of the overarching paradigm for PRS - RV causal opposition with respect to complex traits has been limited. Importantly, prior methods investigating the relationship have been non-parametric. These non-parametric methods do not differentiate main effects and interactions, and they lack interval estimation to quantify uncertainty in results. Moreover, the lack of statistical formalism has hampered clarity on the method and extensions to address study design as well as confounding by ancestry and alternative genetic models.

We develop a formal statistical approach to genetic heterogeneity which we call the Causal Pivot (CP). The method can be applied in a cases-only, controls-only or a case-control design; here we focus on the cases-only application. The CP is based on a structural model of disease (**Figure 1**) as well as the well-known collider bias concept^25–27^ in which conditioning on an outcome induces a synthetic correlation between otherwise independent causes. Under these conditions, if we observe individuals’ disease status and condition on their outcome – either case or control —then, within those outcome groups, the independent causal factors will have induced correlation in well-defined patterns. Importantly, model factors will show no induced correlation if the factors are unconditionally independent and not mutually causal. We view outcome-induced association between causal factors as a feature to resolve heterogeneity and to drive discovery instead of as a source of bias in the research process.^28–30^

**Figure 1.**
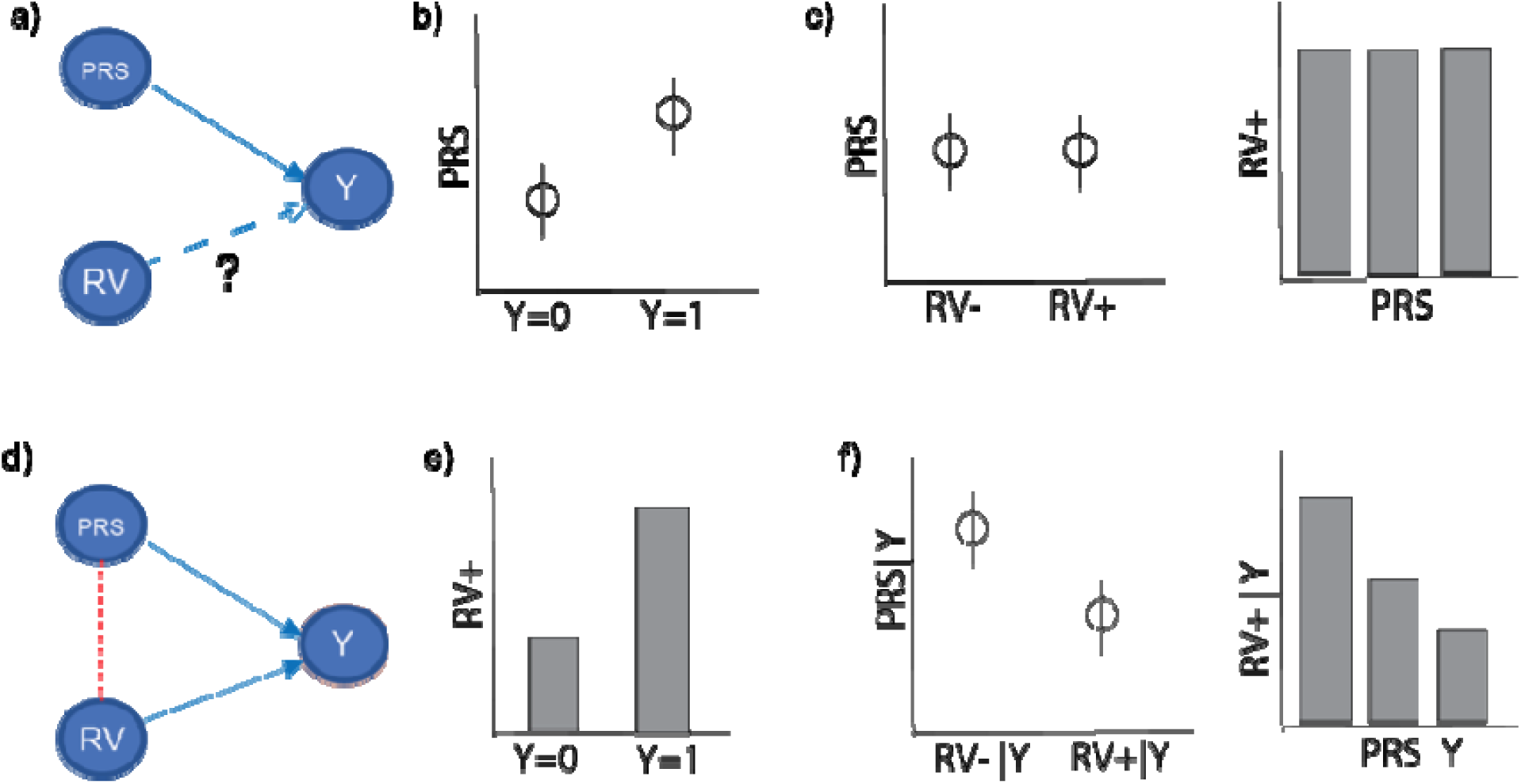
Core Logic of the Causal Pivot. **Panel a:** Basic model. In the base model PRS is a known causal factor (solid line and arrow) terminating at Y; RV is an independent factor whose effect on Y is unknown (dashed line) and being evaluated. **Panel b:** Causal impact of the PRS on Y. PRS is correlated with Y manifesting as a higher PRS in cases where Y=1 than in controls where Y=0. **Panel c:** Model of unconditional independence between PRS and RV. Unconditional independence results in a lack of association between PRS and RV: there is no correlation in the frequency of the RV+ with the PRS tertiles. **Panel d:** RV influences phenotype Y. The solid arrow indicates a causal effect of RV on Y. The red box around Y indicates conditioning on the phenotypic state, such as selection of the Y=1 subpopulation, leading to an induced correlation between PRS and RV. This collider-induced correlation -- dashed red line -- connects PRS and RV. **Panel e:** The frequency change in RV+ when RV is a cause of Y. **Panel f:** The collider-induced correlation between PRS and RV+ conditional on Y. This correlation manifests as a Y-conditional shift in PRS between RV+ and RV−; alternatively, the rate of the RV+ state covaries with PRS conditional on Y. This collider induced correlation between PRS and RV juxtaposes with the unconditional independence of the PRS and RV (**panel c**). The Causal Pivot recruits the collider induced correlation together with the rate change in the RV (**panel d**) as signal to infer the RV−Y relationship.

The CP is a highly general statistical framework built using the tools of SCM and Bayes rule. This approach leads naturally to a likelihood framework, maximum likelihood estimation, confidence intervals, and to a CP likelihood ratio test (CP-LRT). The same mathematical structure applies in the context of both binary and quantitative traits. We perform power analyses and compare the CP-LRT to alternatives ^20,21^. Importantly, the CP-LRT derives power from both the rate change in RV given disease as well as the conditionally induced dependency between the causes; alternative methods may fail to recruit both sources of information. This insight leads to demonstration of the robust properties of the CP-LRT in terms of sensitivity and specificity.

Using human data from UKB we demonstrate the performance of CP-LRT in demonstration examples of breast cancer (BC), hypercholesterolemia (HC) and Parkinson’s disease (PD); we focus on the respective known disease genes *BRCA1*, *LDLR* and *GBA.* The CP procedure successfully identifies the relationships of these known disease genes to their respective diseases. We then extend CP to address confounded structures focusing on ancestry. We successfully demonstrate two approaches to address ancestry confounding using matching and inverse probability weighting. Finally, we develop a pathway burden test to demonstrate the utility of cases-only CP to investigate oligogenic models for pathway contribution to complex disease using lysosomal storage in PD. Our results demonstrate the generality and potential of CP as an approach to causal discovery and to address heterogeneity in complex disease genetics.

## Methods

### Causal Models and Graphs

Structural modeling is also called causal analysis or structural causal modeling. Our methods are developed through structural causal modeling (SCM). In the structural framework directed acyclic graphs are used to express model assumptions where directed arrows depict stochastic dependencies and the lack of a directed path between nodes indicates unconditional independence.

We consider simple models with three or four nodes and two or four edges, respectively (**Figure 1 a, d** and **Figure 4 a,b,c**). The disease outcome variable is denoted **Y**. This outcome represents the phenotypic status, for example as recorded in the UKB metadata by ICD-10 codes or biomarker measurement information. This outcome variable can be binary or continuous. Initially we focus on two genetic exposures: a PRS -- a continuous variable -- which we will denote as **X** representing the combined causal contribution of common variation (MAF >0.01), and a binary RV state variable denoted as **G** that represents the presence of a strongly acting RV. This binary RV represents the state that an individual is positive for any of an ensemble of strongly acting RV that in themselves may be vanishingly rare or singleton but in aggregate can have frequency approaching 0.001 or larger in a population or study sample. These **X** and **G** are depicted as nodes with edges incident to the **Y** outcome (**Figure 1 a,d**). Later we extend this three-node model to a four-node graph (**Figure 4 a,b,c**) where the PRS and RV variables are coordinately influenced by ancestry denoted as **A.**

#### Causal Pivot

When two causes are incident to the same effect that effect is called a collider for the two causes. Conditioning on the collider outcome induces an observational – but not causal – correlation between the distinct causes. This situation is a well-known source of bias in epidemiologic studies because researchers may unknowingly condition on the outcome and report the induced association between variables **X** and **G** as causal between these variables.

The CP exploits collider induced correlation as a source of signal rather than noise. As we show, when one causal variable incident to the outcome in a collider structure has a **known effect on the outcome** – such as the effect of the **PRS** on **Y**, then the outcome-conditionally-induced **RV−PRS** correlation can be used to estimate and to test the **Y** relation to the candidate cause, in this case **Y∼RV** relationship. The Null Hypothesis is an absence of the **Y, RV** relationship, and in this situation no induced correlation will appear between **X** and **G** conditional on **Y**. As we show in the **Mathematical Supplement**, the CP is generally applicable and not limited to genetics.

### Conditional Analysis

Here we present an overview of the mathematical derivation of the CP approach; more details are in the **Mathematical Supplement**. In what follows we refer to the variables PRS as **X**, RV as **G** and disease outcome **Y.** The CP derives from manipulation of the probabilistic factorization determined by the graphical model (**Figure 1a**). The joint density under the model is:

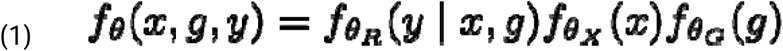

In the expression, represents the respective probability density functions for their arguments, and the bar symbol expresses conditioning on variables to the right-hand side of the bar. The parameter vector θ represents the unknown parameters governing the probabilistic structure of the system; for clarity we emphasize that the parameter θ is comprised of the parameters θ_R_ that determine the forward conditional relationship between **Y** and **X**,**G** – also known as the outcome model; the separate parameters θ_XG_ determine the distributions of **X** and **G** – which in the simple model are exogenous and independent and can be represented separately as θ_X_ and θ_G_.

The Causal Pivot procedure follows from application of Bayes’ rule to the probabilistic factorization implied by the graphical model. In notation, application of Bayes rule and independence of **X**, **G** leads in (1) leads to:

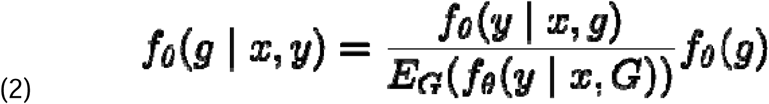

For **G** representing RV status, the expression (2) states that the conditional probability for a sampled individual to bear an RV – denoted by **G=1** – or to not have an RV -- denoted **G**=0 – when **X**, **Y** are conditioned upon is given by the ratio of the forward conditional probability of their phenotype **Y=y** given PRS **X=x** and **G=g** divided by the average probability of **Y** integrating out **G=g** multiplied against the prior probability of **G=g**. This factorization is general and does not depend on the model specification of **f(y | x,g)** or **f(x)** or **f(g)**; moreover, the factorizations in (1) and (2) do not depend on a univariate restriction on **X**, **G** or **Y**, nor does it depend on the binary character of **G** or the distributional character of **X**. The form holds under **Figure 1a**.

### Model Specification

It is useful to provide a parametric model in order to perform analyses, make calculations, enable parametric testing and to perform power comparisons. Non-parametric approaches also apply, but parametric modeling provides both mathematical insight and yields practical results such as confidence intervals. We adopt a generalized linear model framework for the forward regression model of **Y** given **X** and **G.** The general linear model framework encompasses both logistic regression in the case of binary outcomes as well as linear regression for quantitative traits. The general expression for the forward conditional expectation of **Y** given **X** and **G** is:

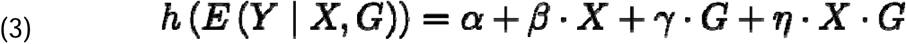

In (3) we do not specify values for the random variables **X**,**G**,**Y**; instead we focus on the structure of the conditional expectation of **Y** as a function of **X**,**G,** Examples for the function h() are the logit or log-odds for binary outcomes, such as for a binary trait like BC; the identity function can be used for continuous **Y**, such as in the case of LDL measurements. We assume the parameters α and β are known or estimable from population data; we also assume knowledge of a baseline frequency of RV, the value ω. In many cases the estimate of the **X**, **Y** relationship is reported. If not, the easiest way to estimate it is to apply a forward model to unconditional **Y** including **X** and exclude **G.** Although the α parameter may depend on **G** we argue that it is negligible because **G** is rare. We adopt this approach to determine α and β in our demonstration work. The frequency of rare variation (ω) is frequently estimable from population databases such as gnomAD^31^ or from UKB summary information.

### Likelihood Analysis Conditional on Y

To examine the properties of the CP we construct procedures for the possible effect of RV using the factorization in (2) to determine a likelihood function for observed data conditional on **Y=a.** We carried out this analysis for both binary **Y** and for Y>δ for continuous **Y**. In the case of discrete Y, we have:

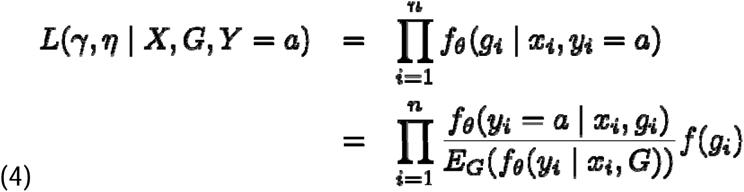

Taking logarithms and emphasizing that the unknown elements of θ τηατ concern the impact of the RV on outcome are γ,η:

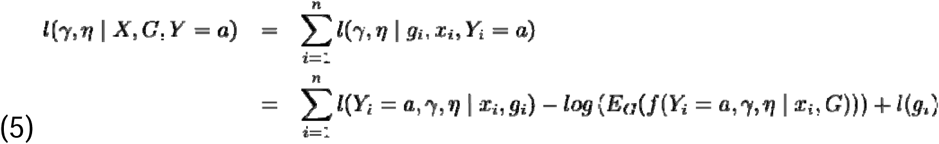

As shown in the **Mathematical Supplement** this likelihood function can be maximized in the variables γ and η to produce maximum likelihood estimates; this analysis is performed by differentiating the log-likelihood with respect to γ and η and solving for the root where the derivative is zero; a check can be performed to ensure the second derivative is negative at the root so that the point is a maximizer. Confidence intervals may be obtained using the observed estimate of the Fisher Information using the empirical means of the squared derivatives of the log-likelihood (see Supplemental). We performed this analysis for both the logistic and the liability model for a continuous trait. For the continuous trait we consider the circumstance where the observable sample comprises individuals with a sufficiently extreme quantitative trait Y>δ. Ιndividuals in this class are deemed cases and individuals not in the class are deemed controls; we suppose we have access to the trait values.

#### Likelihood Ratio Test

As mentioned, the likelihood approach determines maximum likelihood estimates (MLE) by solving for the roots of the derivatives of log-likelihood equations treating these as functions of the unknown parameters. A likelihood ratio test can be constructed by plugging the maximum likelihood estimates into the log-likelihood function to compute a test statistic. The Null Hypothesis is that the parameters γ and η are 0. If the MLEs for γ, and η are substituted into the likelihood function, then under the Null Hypothesis the asymptotic distribution of: −2 * l(0,0 |X,Y,G) - l(γ, η|X,Y,G) is a chi-square on two degrees of freedom. The procedure considers the influence of the RV – both individually and in its potential interaction with the PRS. Additional details are provided in the Mathematical Supplement.

### Logistic Model

The logistic model considers the disease outcome to be binary where the log-odds of disease is linear in the explanatory variable. We additionally treat RV as a binary variable. In the initial treatment there is no additional stochasticity in the conditional distribution of the outcome **Y** given **X** and **G** except for the coin-flipping variability of a Bernoulli trial; such extensions using random effects and the frailty model are possible but out of scope in this paper. The log-likelihood analysis conditional on **Y=a** produces four different sums: the sum of contributions for **Y=0** where **RV=1**, **Y=0** where **RV=0**, **Y=1** where **RV=1**, and **Y=1** where **RV=0**. Breaking the likelihood calculation into these four parts is both computationally practical and conceptually useful. In cases-only analysis, there is no contribution where **Y=0.** The details of the log-likelihood analysis and derivative derivation appear in the Mathematical supplement.

### Liability Model

The Causal Pivot liability model considers conditioning on the outcome Y>δ for a quantitative trait that has a linear outcome model as in (3) where h() is the identity function and where outcomes have additional mean zero Gaussian noise. The approach generalizes to non-Gaussian noise, but the Gaussian situation is both canonical and illustrative. Here the model considers the outcome variable Y | Y>δ, x, g. Introducing an indicator variable for the conditional state, the forward model has a conditional density when Y>δ:

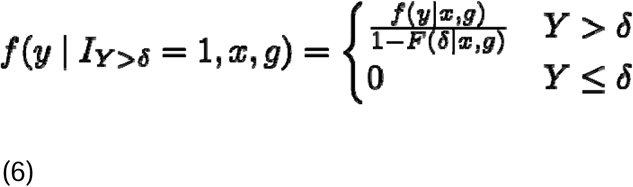

For situations where the outcome value is less than δ we have:

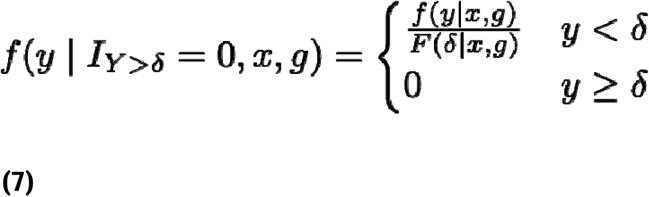

To perform analysis with this liability model we use the general expression in (2) that becomes expression (5) after applying the log-likelihood transformation. As with the logistic model there are four configurations for binary RV: where the outcome Y< δ and G=1, where Y<δ and G=0, where the Y>δ and G=1, and where Y>δ and G=0.

### Power Analysis

To perform analyses, we built a forward simulation procedure to generate outcome data under both the logistic and liability frameworks. For each scenario, we simulated RV as a binary variable with frequency 0.001. For PRS we generated unit normal outcomes. We fixed β according to values consistent with real data. We let γ and η range across many possible values. For logistic regression we set α=−2.2 and β =0.5, we set γ and η to a collection of alternative values. We generated **Y** outcomes according to the inverse transformed log-odds. We then applied the likelihood expression in (5) and solved for the root of the derivatives of the log-likelihood by numerical procedure to obtain the MLE. We then computed the likelihood ratio test statistic. We performed 1000 simulations at each model parameterization, and we recorded outcomes. To determine power, we calculated the proportion of simulation runs where the observed p-value was less than 0.05. For the liability model we generated outcome variables in a parallel fashion, first generating X ∼N(0,1) and G∼ber(0.001) and fixing the parameters α, β and a grid of alternative **RV** effect sizes. We set δ to be the 92%-ile of the trait distribution.

### Cases Only Analysis vs Case Control Conditional Analysis

Our CP procedure can be applied to cases-only data, to controls only data or to case-control data taken together. In all of these situations, data analysis proceeds according to a conditional analysis using the fundamental factorization in (2) and to application of the maximum likelihood procedure in (5) followed by a likelihood ratio test. We focused on comparison of cases-only CP analysis against the traditional forward regression approach and alternative cases-only analyses.

### Comparator Tests and Robustness Analyses

The CP-LRT benefits from the full information in the stochastic system as described by the structural causal graph and as encapsulated in the conditional likelihood function. This conditional likelihood gives the expected rate change in **G** given both **X** and **Y**. We consider two comparator non-parametric models to our CP-LRT. First, we consider a Z-test procedure where we assume the frequency of RV status in the source population – deemed ω -- is exactly known. Under the Null Hypothesis where RV status has no effect on outcome, the expected number of RV in the outcome group **Y=a** would be binomial with expectation N_Y=a_ω and variance N_Y=a_ω(1-ω). We take **Z** to be the difference between the observed RV+ disease individuals (N_Y=a,RV+_) and the expected under H_0_ standardizing by square root of variance. This comparator focuses on the rate change of a functional variant state in the disease population and ignores the PRS. Alternatively, prior work^20^ proposed a Wilcoxon rank sum test (also known as Mann-Whitney U-Test) on the PRS values among RV+ and RV− individuals conditional on **Y=a**. The Wilcoxon is a non-parametric test for a shift in means between two populations, in this case RV+ and RV− conditional on **Y=a** or **Y>**δ. In contrast to the alternatives, the Wilcoxon test does not utilize prior knowledge of RV frequency information.

### Robustness Analyses

Using simulation we compared the Type 1 error of the CP-LRT and Z-test when **G** has no effect and scrutinize the impact of under-specification of the rate ω. The type 1 error is the rate at which the test falsely identifies an effect of **G** when there is none. These analyses were performed across a range of under-specification of the RV rate which we deem ω*; the true simulation parameter values in these analyses are α=−2.2, β=0.6,ω=0.001, γ=0,η=0.

We then consider the power (1-Type 2 error) impact of over-specifying the RV rate. We examine this situation in the context where α=−2.2, β=0.6, ω=0.001, γ=2.2, η=0. In this case, there is an RV effect, but oversetting the value of ω* makes it more difficult for the statistical procedures to detect. We examine a range of values ω* up to and beyond the expected conditional frequency of RV.

### Impact of PRS Effect Size

We examined the impact of the known causal factor’s effect strength on CP performance. We focused on the binary trait outcome context, and we evaluated the impact of PRS effect by letting the β parameter vary. We considered both the impact on the conditional mean of PRS given RV and trait outcome Y as well as the conditional mean of the RV state – which equates to the probability of RV+. The conditional mean is derived by taking the expectation with respect to the conditional density as determined by the factorization (2). We consider the Odds Ratio (OR) for RV+ conditional on Y and set the PRS value X=−1 and X=1. For the conditional mean of PRS given RV and Y, we note there are 4 conditional mean functions to consider conditioning on states of trait Y and RV denoted G: Y=1, G=1; Y=1, G=0; Y=0,G=1;Y=0,G=0. We also evaluated the impact of PRS effect size on statistical power. To emphasize the role of induced correlation we misspecified the RV rate.

### UKB Data

We analyzed data drawn from the UKB under an approved application (Project 98786) titled “Genetic Heterogeneity in Diseases with Complex Inheritance”. We used hypercholesterolemia (LDL direct | Instance 0 ≥ 4.9 mmol/L; Data-Field 30780), breast cancer (ICD-10 C50; Data-Field 40006), and Parkinson’s Disease (ICD-10 G20; Data-Field 131022) as disease models with expected positive findings. Cases were filtered based on inferred European ancestry, availability of exome and genotyping data, female-only for breast cancer, and availability of a phenotype-specific polygenic risk score. The final samples included: hypercholesterolemia (HC cases 24656; controls 347889); breast cancer (BC, cases 12479; controls 198311); and Parkinson’s (PD, cases 2949; controls 388198). As this study exclusively utilized de-identified data from the UKB, it qualifies for exemption from ethical review under the guidelines for research involving non-identifiable human data. Cases where an individual had more than one of these disease phenotypes were excluded.

### PRS

We used PRS made available through the UKB (Field ID 26220 Standard PRS for BC; 26250 Standard PRS for low density lipoprotein cholesterol; and 26260 Standard PRS for PD).^32^

### Exome Variant Extraction and Classification

We used PLINK2 to read genotypes from the UKB exome BGEN files (Data Field 23159) and extract variants within our target gene lists (see bedfile https://github.com/chadashaw/causal-pivot/) for the disease-defined cohort samples. The extracted genotype files were converted to Matrix Market format using a custom python script. Variants were then annotated with OpenCRAVAT; annotators included those pulled from the OpenCRAVAT respository (gnomad3, revel, cadd_exome) as well as custom annotators (tcg_clinvar, AlphaMissense, ESM1b) produced from publicly available data. We exclude variants that were absent (homozygous-ref) across all samples.

We focused on three example genes for the demonstration diseases: *BRCA1* for BC, *LDLR* for HC and *GBA* for PD. We used the OpenCRAVAT annotations to classify rare deleterious and/or pathogenic variants within these 3 genes. Pathogenic variants were ClinVar “Pathogenic” or “Likely Pathogenic” without conflicting or qualifying annotations. Additionally, we included rare stop-gain or frameshift variants that were not found in ClinVar and had a gnomAD v3 Non-Finnish European allele frequency of less than 0.1%. We evaluated as RV+ those individuals with at least one alternate allele for any rare LoF or ClinVar pathogenic variant within each gene. To ensure independence between our two causal factors (RV status and PRS) we exclude all variants from each gene strongly associated with the PRS for that gene’s demonstration disease. We use logistic regression to estimate the outcome unconditional statistical significance of any correlation between individual variants and their associated PRS and remove those with an effect p-value of less than 0.05.

We extracted relevant cohort metadata (age, sex, disease indicators, LDL, and ancestry components) from the UKBB cohort browser.

### Controlling for Ancestry

We considered potential confounding between PRS and RV arising from shared Ancestry, denoted A (**Figure 4**). We adopted two approaches to address this situation: matching and inverse probability weighting. With both methods we focused on the conditional outcome where **Y=1**, cases only analysis. We expect that if RV increases the risk that **Y=1** then a negative association will appear between RV and PRS such that affected individuals (**Y**=1) with RV+ will have an average lower PRS than affected individuals with RV− after conditioning on **A**. To perform the matching, samples in each of the three cohort groups were differentiated based on positive disease (case) status. We determine a distance matrix between all samples using the ‘distances’ package in R. In the absence of eigenvalues for the available PCs in UKB we employ an exponentially decreasing weighting function (scaled to unity) [decay = e**^r^**^*5^, weights = decay / sum(decay) **r = 0.637**] to weight squared differences between observations for each PC in descending order, summing these to determine a distance value. RV+ cases were matched to the closest (k=2) RV− cases using these calculated distances. For each RV+ case, the difference between the observed PRS value and the mean PRS value for the matched RV− cases is calculated. An overall test statistic was then calculated by taking the average of these values over all RV+ cases. Permutation testing was performed using the RV+ status as the treatment variable. A 1-tailed permutation test of means was then performed using the observed test-statistic against the permutation distribution.

We also perform an analysis using inverse probability weighting. Briefly, inverse probability weighting is a general method for adjusting outcome data in causal analysis to account for unequal representation of the causal exposure that correlates with an observable common cause. In this situation the outcome variable is taken to be the PRS among the affected individuals (Y=1) and the exposure is the binary RV; the common cause is ancestry. We used the random forest method to perform a machine learning predictive analysis for RV+ using the ancestry principal components as predictors in each disease cohort. The outcome of random forest is a probability of RV=1 for each observation. We then computed the inverse probability weighted sum of PRS among RV+ and subtracted the sum of weighted values among the RV− cases, using one minus the probability of RV+ as the weights of the RV− cases. We used permutation assignment of the RV status to determine a null distribution and a one-sided p-value.

### Test of RV Load in Individuals

We tabulate a load of variants in a biological pathway as an alternative count-valued candidate cause to test using CP. For this oligogenic load analysis we focused on the lysosomal storage pathway in PD. We enumerated qualifying variants (see method above) in 54 lysosomal storage genes that were also present in UKB exomes. We counted the number of variants in each individual. To examine the potential induced association of this count variable with PRS conditional on disease we used Poisson regression with a log-link to consider the average RV count as a rate that depends on PRS. Briefly, in Poisson regression the logarithm of the rate of RV is modeled as a linear function of the PRS. We used the likelihood ratio test for Poisson regression as implemented in the R glm() method to determine p-values.

## Results

### Comparative power analyses

We hypothesized our Causal Pivot LRT (CP-LRT) could have promising statistical properties to discover RV causal candidates by exploiting a PRS as a known cause. The model assumes the known factor PRS is previously established and has a defined effect size. In our power analyses the candidate cause is a binary variable representing the presence of at least one qualifying RV in an individual. This variable is assumed to have a known sparse rate of occurrence in the general population, and we used the value 0.001. Simulation-based power analyses are presented in **Figure 2a** in the context of a binary outcome trait considering the possibility of both a main effect of RV as well as a PRS-RV interaction. In all simulations the forward logistic regression using the full data of both cases and controls had the highest power; this test is using approximately an order of magnitude more subjects than cases-only analyses because n=500,000 for case-control, vs n_c_ of approximately 50,000. Importantly, using only cases – approximately 10% of the population – CP-LRT achieves 90% of the power of the case-control analysis for the CP-LRT when the main effect of the RV is at least as large as the effect of the PRS. In the situation of cases-only analyses with no interaction effect, the Z-test looking for overrepresentation of RV+ individuals had the highest power. When the RV main effect is sufficient and there is an interaction between RV and PRS the CP-LRT had superior power compared to the Z-test. The CP-LRT has superior power over the Wilcoxon test (purple curve) when there is no interaction effect as shown by the CP-LRT (red curve above solid purple line), but the Wilcoxon test has higher power for weaker acting RV when there is an interaction.

**Figure 2.**
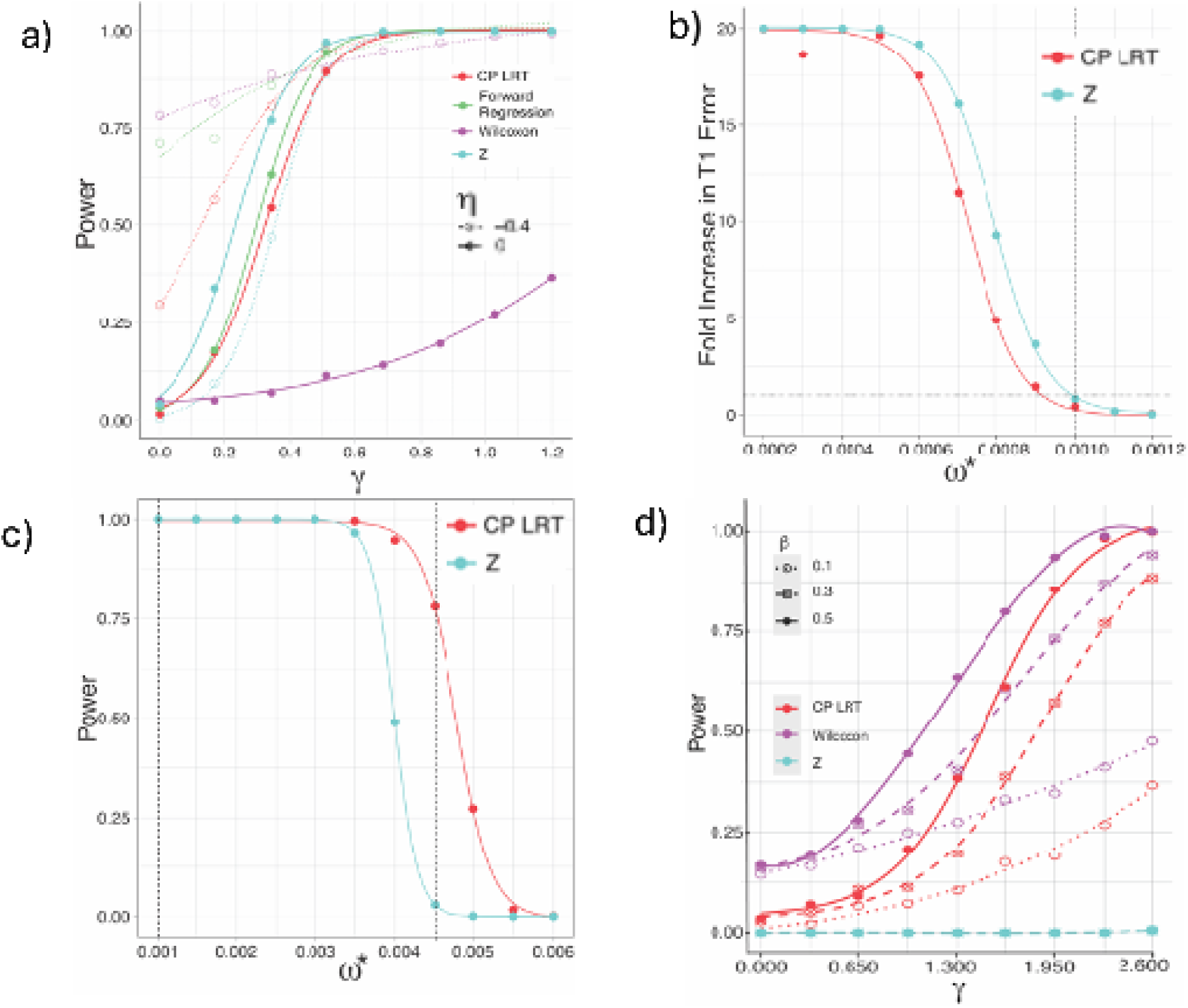
Simulation Analyses. The figure presents simulation analyses comparing the CP-LRT to alternatives focusing on the binary trait context. Monte Carlo methods were used to generate the results in panels a-d; these simulations generated PRS and RV status which were then used to simulate trait outcomes under the corresponding logistic model. Alternative analyses were then performed to detect the effect of the RV and to compare alternative approaches on the same simulated data. Detailed markdown documentation and code to produce these analyses are provided (Supplemental: pivotr.pdf). **Panel a:** Traits were generated using a logistic model with causal effects from PRS and RV status. The simulated population size was n=500,000, comparable to the UK Biobank (UKB) scale. The intercept was set to α=−2.2, corresponding to a disease prevalence of 10%, and the PRS effect was set to β=0.5, consistent with three UKB demonstration diseases. The RV frequency was ω = 0.001. The x-axis represents the main effect of the RV (γ), and the y-axis shows statistical power determined as the proportion of simulations that reject the null hypothesis at a nominal Type I error rate of 0.05 (i.e. p-value<0.05). Power comparisons include the CP-LRT (red), Wilcoxon rank-sum test (purple), Z-test (cyan), and forward logistic regression using both cases and controls (green). Dashed curves indicate a PRS-RV interaction effect (η=−0.4). In the absence of interaction effects, the Wilcoxon test had markedly lower power, and the Z-test outperformed CP-LRT but did not differentiate between main effects and interaction effects nor produce parameter estimates. **Panel b**: Type I error robustness. The y-axis shows the fold increase in Type I error relative to the nominal level of 0.05. Simulations held γ=0, η = 0, and α, β, ω as in **panel a**, while underspecifying the RV frequency (ω*). The CP-LRT (red) retained Type I error robustness with values near 0.05 despite a 10% reduction in ω*, while the Z-test (cyan) experienced a sharp increase in Type I error as ω* decreased. **Panel c**: Sensitivity to RV frequency overspecification. We set α=−2.2, β=0.6, γ=2.2; the x-axis represents RV frequency (ω*) used in analysis, exceeding the true frequency (ω = 0.001). Power comparisons between CP-LRT and Z-test show that over-specifying ω* diminishes Z-test power. At ω* = 0.045, the expected RV frequency in the Y = 1 population, the Z-test power approaches 0.05, while CP-LRT maintains ∼75% power by leveraging structurally induced correlation between PRS and RV. **Panel d**: impact of the strength of the PRS. The x-axis ranges the RV effect size γ the other parameters are fixed (α=−2.2,η=−0.4, ω=0.001); ω* is misspecified to its empircal frequency in the disease population. The Z-test has no power in this context. Both the Wilcoxon and CP-LRT show power increase as the strength of the PRS increases. The Wilcoxon has superior power but does not distinguish the main effect from interaction. Results for the liability model are included in the **Supplmentary Materials**.

We also performed robustness analysis focusing on the binary trait context in cases-only analysis. In this situation both the Z-test and the CP-LRT rely on prior information concerning RV frequency. Therefore, we explore the impact of errors in the specification of this frequency. **Figure 2b** considers the situation where there is no RV effect, with both γ=0 and η=0; the unconditional RV+ rate is incorrectly specified lower than its true value across a decreasing range. Results show that the CP-LRT had lower Type 1 error rate compared to the Z-test, as indicated by the cyan Z-test curve dominating the red CP-LRT curve. The CP-LRT retained a type 1 error of 0.05 when the RV rate is misspecified by 10%.

Conversely, the rate parameter for the RV might be falsely specified above its true value. Figure **2c** shows this situation when the RV effect is set to α=−2.2, with γ=2.2 and the PRS parameter is β=0.6 – values consistent with the UKB analyses for BC. In this situation, the CP-LRT had higher power to detect an effect while the Z-test power is sharply reduced, as indicated by the red CP-LRT curve dominating the Z-test cyan curve. Interestingly, the CP-LRT retained a power of ∼75% when the RV frequency is set to the expected RV frequency among cases of approximately 0.0045; in this circumstance the Z-test power is the nominal error rate of 0.05.

In **Figure 2d** we explore the impact of the strength of the PRS on power for the alternative approaches; this was done by varying the parameter β and looking at power as RV effect changes. As in **Figure 2c** we set ω to its value in the affected population to focus on the induced correlation between PRS and RV as PRS strength varies. In this case the Z-test has no power as shown. We ranged the RV effect size and evaluated at three choices of β. In all analyses power increases as β increases. Results show that in this case of misspecification and variation in β the Wilcoxon test has highest power. The CP-LRT has power greater than 0.7 when β=0.5 and γ>1.5.

In addition to power analysis we performed investigations concerning the impact of PRS explanatory power on the conditional mean of PRS given RV status and outcome as well as the conditional probability of RV+ given PRS and outcome. These results are presented in the **Supplemental Materials Figure 1 and Figure 2**. The results show that as the PRS effect size increases there is a strongly increasing impact on the odds of RV+ status among individuals with low PRS scores. The conditional mean of the PRS is also influenced by increasing the effect size of the PRS; interestingly, the conditional mean is more differential between RV+ and RV− among controls rather than cases. The differential between RV+ and RV− in the conditional PRS mean among the controls is increasing in the PRS effect size (**Supplemental Materials Figure 2**).

For the liability model (**Supplemental Materials Figure 3**) we simulated according to a linear model for continuous quantitative outcomes. We modeled our analyses on the example with parameters α=0, β=0.3, ω=0.001, and we used a cutoff value of δ=1.13, corresponding to the 92%-ile of the trait distribution, consistent with clinically relevant LDL cholesterol cutoffs. Conditional cases-only analysis corresponds to the situation where observed samples are restricted to those with trait values over a cutoff. In this situation, the CP-LRT has greater power than the Wilcoxon test when there is no interaction effect. When the interaction effect is strong the Wilcoxon test has power of unity. The forward linear regression model using both case and control data has the strongest power, and the Z-test also has high power.

### Application to UKB data

To assess relevance of CP in real data we used UKB exomes and PRS. We used well studied complex traits with both validated PRS as well as established single gene contributors. The latter allowed us to recruit ClinVar annotations to select known disease-causing rare variants as positive controls. Disease cohorts and variant selection are as described in the Methods. Most variants identified as RV+ were present in only a single individual as shown in **Figure 3 b**. The variants had a range of coding impact as shown in **Supplementary Figure 4.** A detailed enumeration of the variants identified for the CP analysis is included as a Supplement.

**Figure 3.**
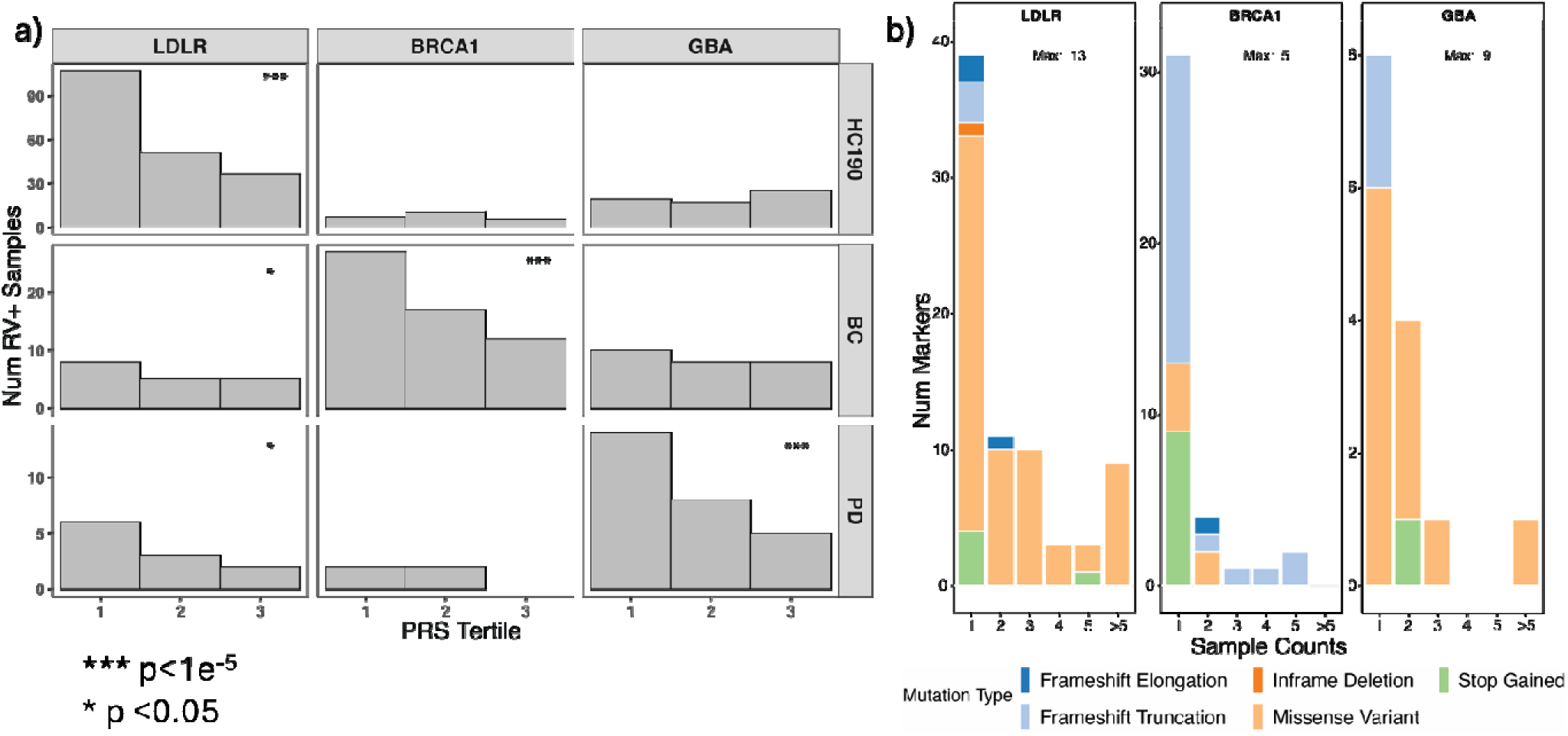
Demonstration of the Causal Pivot in UKB data. **Panel a:** Count of RV+ samples in each PRS tertile. Vertical bars represent the count of RV+ Samples within each PRS-tertile (x-axes). Each row represents a different disease cohort paired with its disease specific PRS – HC, BC, and PD, respectively. Each column presents a different disease gene: *LDLR*, *BRCA1,* and *GBA*. The diagonal of **Panel a** shows the negative correlation between the number of RV+ samples and PRS scores within each PRS-Disease-Gene grouping. CP-LRT was used to determine p-values for all 9 analyses, and the p-values for all tests are highly significant along the diagonal. Off diagonal entries are not significant except for the gene *LDLR* which shows modest association pattern with both BC and PD with p<0.05. More detailed results including confidence intervals for g and h appear in the S**upplemental Table 2**. **Panel b**: Multiplicity and allele type of the RV included in **Panel a**. Most RV occur in only a single sample. A detailed listing of the variants that inform the analysis is provided in the Supplemental materials.

### Cross Cohort Comparisons

We employed a cross-cohort comparative approach to evaluate the CP-LRT. We extracted affected individuals in each disease population and their respective PRS values as well as an RV determination by the presence of at least one qualifying RV (see Methods). For visualization of the relationship of RV+ against PRS we split the respective cohorts into tertiles by their PRS. For each PRS tertile in each cohort we counted the rate of RV+ in each PRS tertile bin; this analysis is repeated for each of the three example genes: *BRCA1*, *LDLR* and *GBA*, respectively. We performed statistical testing by application of our CP-LRT (**Figure 3a**). The three analyses along the diagonal of the graphic are all highly significant (p< 1×10^−6^) indicating that the CP-LRT found an effect in each disease against RV+ status in its respective disease gene. Cross-cohort analyses are mostly not significant, indicating that RV+ status for *BRCA1* and *GBA* are not associated with either PD state or BC state, respectively. The *LDLR* showed a weak but significant CP-LRT association with both BC and PD (**Supplemental Table 1)**. The MLE estimates of the RV and interaction parameters including the point estimates and confidence intervals are provided **Supplemental Table 2**. The point estimates computed using the CP MLE strongly agreed with the results from case-control analyses for both the main effect of RV and the RV:PRS interaction effect (**Supplemental Figure 6**).

### Controlling for the effect of Ancestry

Ancestry may confound the PRS-RV relationship by acting as a common cause for an individual to have a functional RV as well as a high or low PRS. We call this scenario the diamond graph (**Figure 4 a-c**). We used both matching and inverse probability weighting to examine for an outcome-induced association signal between RV and PRS while controlling for this possible ancestry confounding effect. For matching analyses we identified ancestry neighborhoods of affected individuals, and we took the difference of the PRS between matched RV+ and RV− affected individuals. We averaged these PRS differences. The results show that RV+ affected individuals had significantly lower PRS scores on average than ancestry matched RV− individuals (**Figure 4d**).

**Figure 4.**
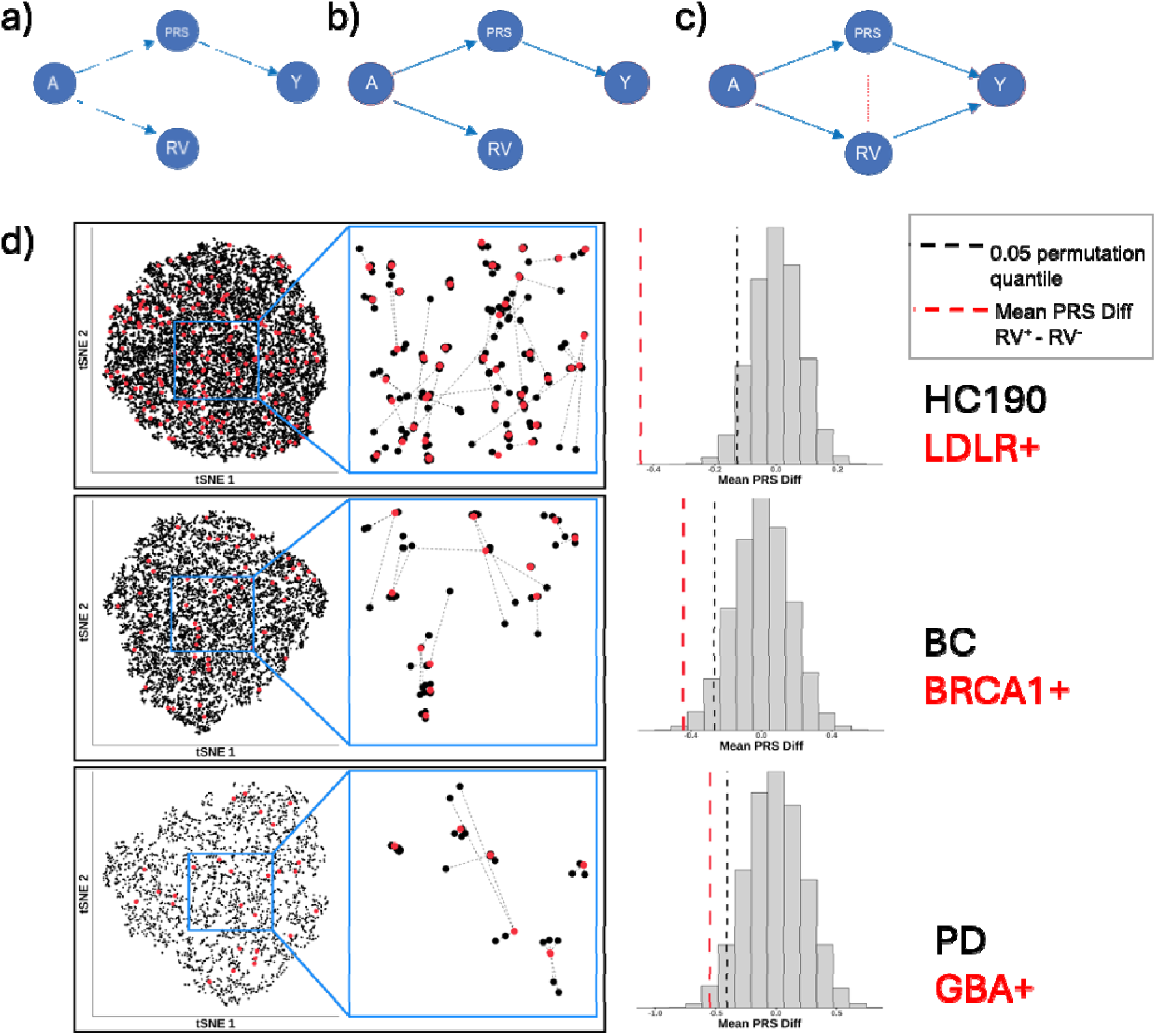
Controlling for Ancestry. Ancestry is a potential common cause – a source of confounding bias – not only between genetic variants and phenotype, but also between alternative genetic causes. **Panel a:** Model scenario where ancestry represented by node A drives both PRS and RV state. **Panel b:** Model in which RV is not a cause of Y. Conditioning on both A and Y there is no induced or residual correlation between PRS and RV. **Panel c:** RV drives Y, and conditioning on both A and Y yields an induced correlation between PRS and RV. **Panel d:** Ancestry matching. Each row represents an example phenotype: HC, BC, and PD, respectively. The scatter plots show RV^−^ (black points) and RV^+^ (red points) cases in ancestry PC space. The scatter plot representation is determined by the first 5 PCs reduced to two dimensions using t-distributed stochastic neighbor embedding (t-SNE). A zoom in view of each row is indicated by the excerpted rectangular region. Each RV^+^ case is matched to the nearest (k=2) RV^−^ cases in the 5-dimensional ancestry space. The mean difference in the PRS of each RV+ and its ancestry matched neighbors and the average of these PRS differences was calculated (dashed vertical red line). Permutation distributions (plotted in the histograms) were generated by randomly permuting the RV states of each sample in the cohort while fixing the ancestry data; the procedure of identifying neighbors and calculating the mean difference in PRS was repeated. The loweer 5% threshold value is shown (dashed black lines). The observed value from the true RV+ ancestry matched data is significantly below the 0.05 quantile in all diseases, consistent with the pattern that RV+ cases have lower PRS than their RV− ancestry-matched neighbors. Additional analyses using propensity and IPW are in **Supplementary Figure 9**.

Propensity scoring and inverse probability weighting is an alternative approach to adjust for common cause confounding. We treated RV+ status as a binary exposure, and we constructed random forest models to predict the chance of this RV+ using the ancestry PCs. The random forest models were successfully predictive of RV+ status, and the results again showed that PRS values are significantly lower in RV+ compared to RV− individuals (**Supplemental Figure 9**). High order PCs were the most predictive of RV+ status.

### Generalization to pathway burden scores

We considered the lysosomal storage pathway in the context of PD as a demonstration pathway for oligogenic load analysis using the CP. The lysosomal storage pathway is the system that supports recycling of macromolecules in the brain, and there are many lines of evidence that point to lysosomal storage as a central pathway in neurodegeneration.^33^ We established a set of 54 human lysosomal storage pathway genes (**Supplemental table**). For each individual we tabulated the number of qualifying RV in the lysosomal storage pathway. We then computed the mean PRS of individuals with disease stratifying on the number of pathway RV. The results are presented in **Figure 5a**; Poisson regression based on the CP demonstrated a significant negative trend in PRS with increasing pathway burden of RV among affected individuals. To emphasize the combinatorial sparsity of the RV data we analyzed the count of two-variant combinations among cases **Figure 5b**. No single pair of lysosomal genes accounted for the majority of the observed oliogenic load.

**Figure 5.**
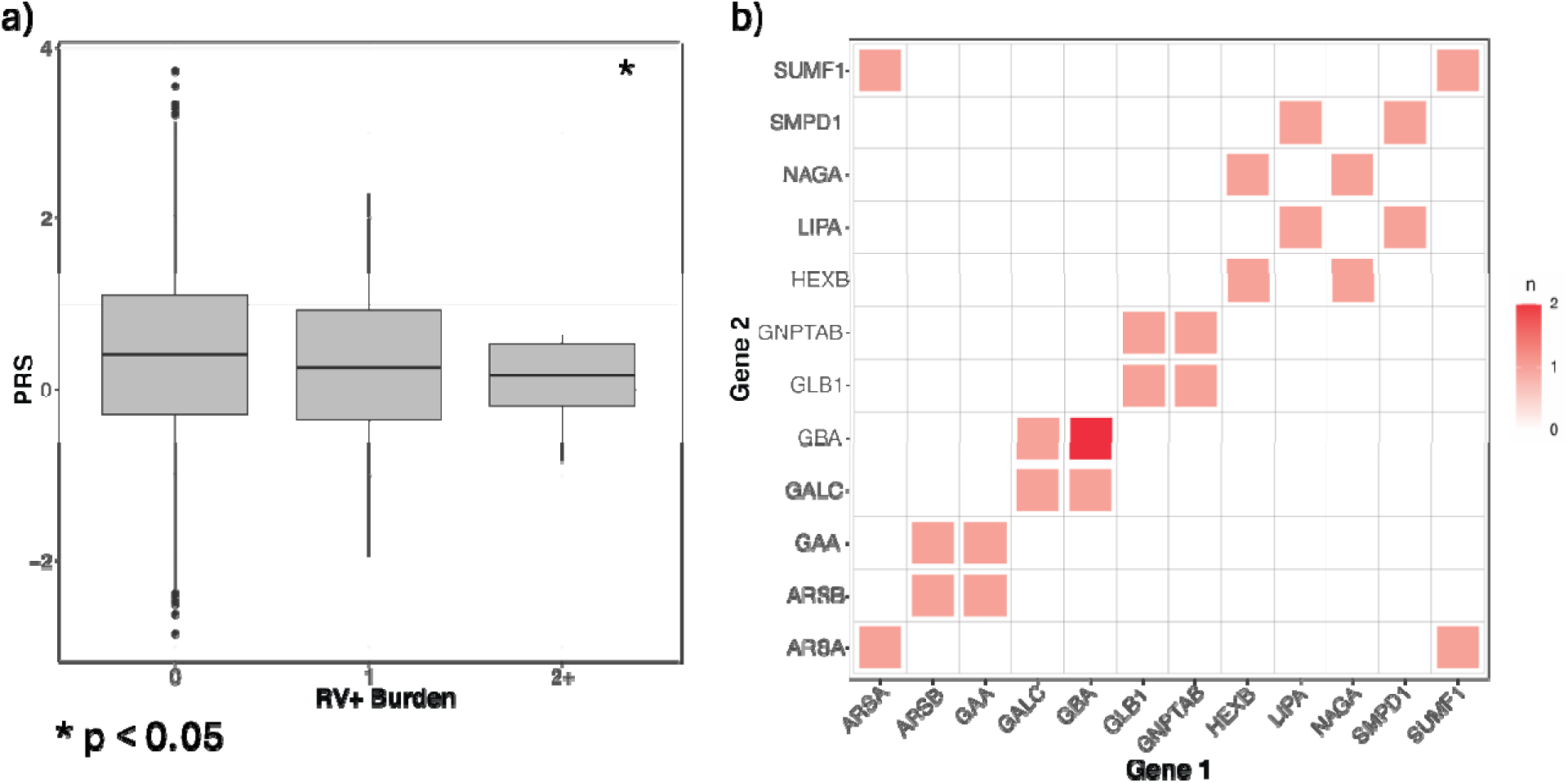
Pathway burden analysis. RVs aggregated by functional pathway can be evaluated as an independent cause by pivot against the PRS, demonstrating that CP-LRT is generalizable to count variables. **Panel a:** shows the PRS values (y-axis) against the count of pathogenic/likely pathogenic RV in the lysosomal storage pathway among PD cases from UKB. Each boxplot presents cases with the tabulated count of RV+ per case. There is a decreasing trend in PRS, and a Poisson regression analysis which models the rate of RV as a function of PRS demonstrates a statistically significant negative association between the rate of RV+ and the PRS value (p=0.03). **Panel b:** Pairwise co-occurrence of RV+ in the lysosomal genes. No single pair accounts for the majority of instances showing the sparse occupancy of positivity for two variants.

## Discussion

To date, the general principles of causal inference and structural causal models (SCM) in complex trait genetics have largely focused on Mendelian randomization (MR)^14,16^ which exploits DNA variants as instrumental variables. MR methods are typically used to investigate mechanisms that flow through a candidate biomarker and are not used to differentiate individuals based on multiple independent causes or causal classes. Given the success of MR, it is likely that SCM can be used more broadly to model genetic effects, suggesting several new avenues for investigating common complex diseases and traits.^34^

To develop the CP we reason that individuals reach the complex disease state through alternative mechanisms that are differentially present or absent among individuals. In most research and diagnostic scenarios investigators have some prior knowledge -- although incomplete -- about causal factors and the strength of their relationship to outcomes. CP explores the utility of induced correlation between a known factor and a candidate cause conditional on outcomes as an additional source of signal for candidate causes. This approach could enable discovery in cases-only analysis because the induced correlation would be present even when appropriate control individuals are not available.

In this study we use polygenic risk scores (PRS)^20,21^ as a proxy for the aggregate genome wide causal contribution of weakly acting common variants to complex disease. This allows us to treat the PRS as a pivot to search for new causes by incorporating induced correlation between alternative disease drivers and PRS by conditioning on outcomes. We note that any known cause, including non-genetic factors, could be either substituted for or added to PRS as a known factor for pivot analysis.

We use simulation-based power analyses for controlled comparison of the CP-LRT approach and alternatives in the cases only context. The Wilcoxon Rank Sum and Z-test are chosen as non-parametric comparators that also apply in cases-only analysis. The Wilcoxon test has been proposed previously^20,21^, and focuses exclusively on the induced correlation between PRS and RV state conditional on disease. Alternatively, the Z-test focuses exclusively on the rate change in RV in the disease population compared to its observed background frequency in the overall population, and it does not use the PRS information. The CP-LRT approach – derived from the structural causal model factorization and Bayes rule -- incorporates *both* the frequency information from the general population and also the PRS-RV conditional dependency.

The results show that in the absence of an interaction effect the CP-LRT has stronger power than the Wilcoxon test, but it is weaker than the Z-test. When there is an interaction effect the power of the CP-LRT exceeds the Z-test, but the Wilcoxon test surpasses it among weakly acting RV with smaller effects. Importantly, the CP likelihood approach results in parameter estimates and confidence intervals; these interval estimates are an important advantage of the procedure over the non-parametric methods which do not quantify uncertainty, produce parameter estimates or distinguish between main effects and PRS:RV interactions.

Robustness analyses for CP-LRT reveal key properties of the method. In the cases-only context, prior information concerning RV frequency is a precondition to apply either the Z-test or the CP-LRT. Importantly, the CP-LRT is informed by the structural model and includes the PRS as part of its procedure, while the Z-test does not. The results show that this difference confers a robustness advantage to the CP-LRT. In terms of specificity, the CP-LRT is more stable against false positives when the population rate of RV is underestimated; the Z-test is highly vulnerable to this under-specification. Conversely, the CP-LRT is also more sensitive when the RV rate is overestimated, and it demonstrates good power when the RV frequency was set to the frequency in the affected population without requiring prior information for RV rate. Taken together the CP-LRT has an advantage in sensitivity and specificity because the procedure is drawing information from the structural model and the conditionally induced correlation.

We also examine the impact of the strength of the PRS on CP performance. We studied the characteristics of the conditional means of both PRS and RV. We see that increasing the PRS effect leads to an increase in the response of the conditional mean and corresponding increase in power; this property is most clearly revealed in the context of misspecification of the RV rate. The Z-test does not incorporate the PRS and is independent of its effect size. The Wilcoxon power also increases with increasing PRS effect. The patterns of the conditional mean functions are consistent with increasing power driving larger magnitude changes in conditional mean. Interestingly, the differential between RV+ and RV− conditional mean of the PRS is larger in controls than in cases. There are circumstances for larger PRS effects where the difference in the conditional PRS mean between RV+ and RV− in cases is zero; this pattern does not hold in controls.

Our analyses of data from the UKB reinforce the validity of the CP model. We chose demonstration diseases with well-established disease genes that are mechanistically associated with their respective complex diseases^35–37^; these diseases also have established PRS with demonstrated risk associations^38–40^. In all cases we observed a strong collider -induced association between the PRS and RV status within diseases and significant CP-LRT results, consistent with our motivating structural model. We also performed parallel analysis using synonymous RV in these same subjects; these synonymous variants do not demonstrate significant collider induced patterns and lack significant CP-LRT findings. Taken together, these results demonstrate results that are consistent with our CP model. Cross disease analyses are also revealing. The induced association with functional RV does not appear between *GBA* variants and BC nor between *BRCA1* variants and PD. Interestingly, *LDLR* variation does show a modest induced correlation effect with both BC and PD. Previous studies have mechanistically implicated cholesterol in diverse disease processes including cancer^41^ and neurodegeneration^42,43^. These findings may warrant further investigation.

An additional contribution of our work is to address ancestry confounding through causal approaches. We extend our graph to consider ancestry as a common cause node for both PRS and RV state. We note that PRS themselves are constructed controlling for ancestry by regression methods^44^, and in our work we avoid RV that demonstrate marginal association with PRS. However, these analyses are limited in power when individual RV are sparse and could miss cryptic dependency. To overcome these limitations we performed matching of RV+ to RV− cases in the space of the global ancestry PCs. We then compared the PRS of RV+ cases to the average PRS of their ancestry matched RV− neighbors. Neighbor matching is a well-established method to non-parametrically correct for complex confounding variables, but this method has not been extensively deployed in genetic epidemiology. We also took a propensity modeling approach. We used machine learning by random forest to predict the probability for an individual to be RV+ based on ancestry PCs. We then used inverse probability weighting using this propensity model to construct a weighted difference of the PRS between RV+ and RV− individuals; this analysis also produced significant results showing lower PRS in RV+ cases, consistent with the CP model for all three example diseases.

Finally, we consider a pathway load as a candidate cause and again utilize the PRS as a pivot. We chose the count of qualifying variants in the lysosomal storage pathway variants in PD because such variation is mechanistically associated with neurodegenerative disease processes both by experimental investigation as well as epidemiologic evidence ^33^. We found significant induced association with the PRS among cases. This finding is encouraging for further investigation. Oligogenic disease models can be difficult to interpret, and the CP framework places them in a causal perspective that reinforces a mechanistic interpretation of statistical findings.

The study of independent causes, in classical genetics called locus and allelic heterogeneity, is central to understanding the genetic architecture of monogenic disorders.^45^ Parallel concepts of causal heterogeneity are needed for complex traits where current methods focus on the average causal contribution of variants rather than their operation in potentially heterogeneous subgroups.^46^ There are already well-known examples in certain complex traits in which the total population of affected individuals includes both those with high polygenic risk along with individuals affected by monogenic forms. In this work, we have developed and evaluated an approach to address this form of causal heterogeneity using a general SCM framework. By examining the possible conditional relationship between a known cause and a distinct candidate cause conditional on the phenotypic outcome we demonstrate that one can establish the candidate’s contribution. Moreover, we show that this approach applies in cases-only, controls-only or combined case-control analyses. The terminology Causal Pivot (CP) is adopted to reinforce that the known cause is used to leverage the conclusion.

The CP is not a foreign concept in medical research. Differential diagnosis is an example of CP thinking. Given that an individual is observed to have some phenotype, when certain causes are ruled out, other causes become more likely. In genomic medicine, the pivot has been observed in somatic cancer, where driver mutations are observed to exclude each other.^47–49^ This exclusion principle is also at the heart of complementation group analysis in experimental genetics. This approach has the goal of deconvolving patient populations into causal groups or subtypes; it has the additional benefit of providing individual level insights following the diagnostic process where evaluating and eliminating alternative potential causes is fundamental to personalized medicine. Our focus in this work is the utility of this approach as a statistical method for discovery in genetic data. In our **Supplemental Figure 1** we show the relevance of our work in a more diagnostic context where we consider the odds ratio of RV+ for individuals with low and high PRS values.

We used simulation and real data to consider CP analyses in a cases-only context. The cases-only test has slightly less power than a comparator case-control analysis, indicating that most of the information about the causal effect of the RV is carried in the cases. It is important to note that such a cases-only analysis is not possible using traditional association tests. The new strategy is therefore ideal when selection on the phenotype is unavoidable. There are many such scenarios relevant to complex diseases. Genotype X environment interaction studies investigate whether the genetic variant and environmental exposure jointly influence the risk, which cannot be tested in those who do not have the outcome. In case-crossover designs the study is restricted to individuals who have already experienced the outcome, and the focus is on understanding the exposure timing in relation to the event within the same individual. Cases-only analysis also applies in the context of somatic mutation outcomes in cancer where analysis is confined to individuals who already have cancer, and the focus is on the correlation between mutations and exposures or germline genetic variation. Another cases only context occurs in drug safety studies that investigate the association between drug exposure specific adverse outcomes; the design also applies to pharmacogenetic studies that assess the drug response among patients who have already experienced a particular outcome, such as an adverse drug reaction. In all these circumstances cases-only genetic analyses are required.

The study of causal heterogeneity in complex disease genetics is a fertile area for research. Significant progress has been made in identifying genes and mechanistic processes that drive complex disease. In this context comparably less effort has been assigned to deconvolving the disease population into subgroups. Our CP and CP-LRT method exploits prior research from GWAS as well as surveys of genetic variation to drive discovery and moves toward assignment of causal types. Additional research in the area of causal heterogeneity using SCM is likely to prove productive.

## Supporting information

MathematicalSupplement

RareVariantTable

CodeSupplement

## Appendices

Supplementary Materials - .pdf

Code Supplement - .pdf

Mathematical Supplment : Derivation of Causal Pivot - .pdf

Table of Annotated Variants - .xlsx

## Acknowledgments

CAS was supported by research funding from the Dan and Jan Duncan Neurological Research Institute. This work was supported, in part, by a grant from Genetics & Genomics Services, Inc as well as funding support from Texas Genomics Consulting.

## Author Contributions

CAS developed the concept of the Causal Pivot and derived the SCM representation. CAS developed the likelihood representation and likelihood ratio test working with TTT. CAS derived the likelihood score equations for the likelihood analysis as well as the ratio representation for simplifying the CP-LRT; he developed the approach to Fisher Information and interval estimation for the CP method. He also developed the matching and inverse probability weighting method for ancestry correction as well as the pathway burden modeling. He supervised the implementation of software and he wrote the manuscript and supplement. *Conceptualization, Formal analysis, Funding acquisition, Methodology, Project administration, Software, Resources, Supervision, Writing – original draft, Writing – review & editing*

*CJW* participated in data curation as well as refinement and execution in software implementation of both power simulations and UKB validations including implementation of the MLE and CP-LRT. CJW also executed the implementation of matching and random forest analysis to control for ancestry. He generated the results and figures for the manuscript. *Data curation, Software, Validation, Visualization, Writing – original draft*

*JWB* worked together with CAS to develop the concept of CP for complex disease and its context in genetic epidemiology. He guided the choice of example diseases and the selection of the proof of concept genes. He contributed to the derivation of the log-likelihood score equations. He provided funding support, and he was strongly involved in writing the manuscript. *Conceptualization, Data curation, Funding acquisition, Project administration, Software, Resources, Writing – original draft, Writing – review & editing*

*JS* contributed to development of the CP concept in neurodegenerative disease and identified *GBA* as a candidate gene for CP analysis in PD. He reviewed and assisted with the manuscript and provided funding support. *Conceptualization, Data curation, Funding acquisition, Writing – review & editing*

*TTT* worked with CAS on development of the LRT for CP and on initial simulation studies. He participated in development of the manuscript. *Formal analysis, Methodology*,

*ND* assisted with implementation of the log-likelihood score equations for both the logistic and liability model. He performed simulation studies for power analysis. *Formal analysis, Methodology*

## Web Resources

*N/A*

## Data and code availability

Code is included in the code supplement.

