## Supplementary material for "The Causal Pivot: A Structural Approach to Genetic Heterogeneity and Variant Discovery in Complex Diseases": MathematicalSupplement

### Mathematical Supplement

#### Causal Pivot

The Causal Pivot is a conditional approach to regression analysis when considering a structural model with an outcome variable $Y$ that is known to depend on one covariate $X$ with a known effect and for which we are trying to discover if $Y$ depends on another separate covariate or covariates which we will generically denote as $\mathbf{G}$. The central concept of the Causal Pivot is to observe or condition analysis of a sample on the outcome $Y$. We then approach the analysis of the relationship between $\mathbf{G}$ and $Y$ conditional on $Y$; we use the structural model hypothesis and the known $Y \sim X$ relationship as part of the signal. By signal we mean that we approach the inference problem for the $Y \sim G$ relationship by considering the conditional distribution of $G \mid X, Y$. In the analysis below we consider $X \perp G$ ; that is unconditionally, $X$ is independent of $G$. Under this assumption, by the collider principle there will be an induced dependency $X \sim G \mid Y$ if and only if $Y$ is dependent on $G$. As we show this induced dependency contains information to contribute to inference of the $Y \sim G$ relationship. The motivating graphical model DAG is:

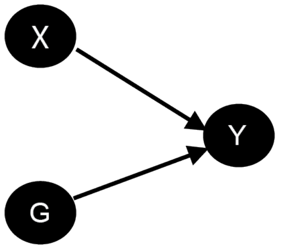


##### Binary G

The simplest case to consider is where $G$ is a univariate binary variable, however the analysis below is not restricted to this case. Without loss of generality we consider a general linear model (glm) framework to describe the dependency of $Y$ on $X, G$. Under this framework the conditional expectation for the outcome variable $Y$ has the form $h(E(Y \mid X, G)) = \alpha + \beta X + \gamma G + \eta X \cdot G$, so that the transformed conditonal expectation of $Y$ is linear in $X, G$ and their interaction $X \cdot G$. Common choices of $h(\cdot)$ are the logistic transform for binary outcomes and the identity function for the standard linear model. We call the glm specification of the forward regression model or the forward model for simplicity. The null hypothesis $H_0$ is that $\gamma = 0, \eta = 0$, so that $Y$ does not depend on $G$. The parameter estimation problem in terms of the forward regression concerns the sub-parameter vector $\theta_{\mathbf{R}} = (\gamma, \eta)^t$ treating the parameters $\alpha, \beta$ as known. The parameters for the distributions of $X, G$ are represented by $\theta_X$ and $\theta_G$,taken together as $\theta_{X,G}$. For the case we will treat in detail below we let $G$ is binary with rate parameter $\omega$ and we let $X$ be Gaussian with $X \sim N(\mu_x, \sigma_x)$ then $\theta_{X,G} = (\omega, \mu_x, \sigma_x)^t$. The key concept is that $\theta = (\theta_R, \theta_{X,G})^t$ where $\theta_R$ has to do with the forward regression part of the problem and $\theta_{X,G}$ has to do with the exogenous casual variables $X, G$.The parameters for $X, G$ do not play a role in the estimation of $\theta_R$ in the ordinary forward regression approach where analysis does not condition or restrict to outcomes of $Y$. We will show that $\theta_R$ and $\theta_{X,G}$ become entangled in the Causal Pivot analysis.

To understand the estimation and inference problem we use the DAG model factorization theorem and Bayes' rule to determine that:

$$f(g \mid x, y) = \frac{f(y \mid x, g)}{E_G\left(f(y \mid x, G)\right)} f(g)$$

when $G$ is binary the expectation in the denominator $E_G\left(f(y \mid x, G)\right)$ takes the form : $E_G\left(f(y \mid x, G)\right) = P(G = 1) \cdot f(y \mid x, 1) + (1 - P(G = 1)) \cdot f(y \mid x, 0)$. This factorization does not depend on the model specifications for $X, G, Y$, it merely assumes that the requisite density functions exist. Note we have suppresed the parameterization in the expression above for simplicity. In the case where $Y$ is also binary and where $A = \{1\}$ we have the simplification:

$$f(g \mid x, 1) = \frac{f(1 \mid x, g)}{E_G\left(f(1 \mid x, G)\right)} f(g)$$

To expose the generality of this approach, consider a discrete or non-negative integer valued $G$ and binary $Y$ where the conditioning set is $Y = 1$,the expectation in the demoninator is $\sum_{G=g} P(G = g) f(1 \mid G = g, x)$, where $f(1 \mid x, g)$ is the conditional density determined by the forward regression model for the outcome variable $Y = y$.

Our goal is decide if $H_0 : f_\theta\left(g \mid x, y\right) = f_{\theta_G}(g)$ is plausible given the data . We note that $f_\theta(y \mid x, g) = f_{\theta_R}(y \mid x, g)$ and that $f_{\theta_G}(g)$ and $f_{\theta_X}(x)$ are $f_{\theta_{X,G}}(g)$ and $f_{\theta_{X,G}}(x)$ respectively. Therefore, we can re-write:

$$f_\theta(g \mid x, y) = \frac{f_{\theta_R}(y \mid x, g)}{E_{\theta_G}\left(f(y \mid x, G)\right)} f_{\theta_G}(g)$$

In the conditional sample situation where $Y \in A$ such as $Y = 1$ the expression is

$$f_\theta(g \mid x, 1) = \frac{f_{\theta_R}(1 \mid x, g)}{E_{\theta_G}\left(f(y \mid x, G)\right)} f_{\theta_G}(g)$$

Importantly, the denominator expectation requires the quantity $E_{\theta_G}\left(s(G)\right)$ where $s()$ is the forward regression density function treating $G$ as random. This expectation depends on the parameter space including the forward regression parameters $\theta_R$ and the exogenous $\theta_G$. In this way we can see that the analysis of $G$ conditional on $X, Y$ has entangled in the conditional density the regression parameters for the forward model with the parameters of the exogenous variables.

**Logistic Model**

For maximum likelihood analysis and for likelihood ratio tests, the likelihood, log-likelihood and derivatives of the log-likelihood with respect to the parameters of interest are the crucial. In the tradition of likelihood analysis we consider the likelihood to be a function of the parameters treating the data as 'fixed' or the conditioning value. A challenging element in this case is that the data is reconsidered in terms of conditioning or restricting to y-values that are observed; we are refactoring the stochastic model so that parameter estimation is considered in the conditional case. The data could be restricted to $Y = 1$, $Y = 0$ or it can be pursued with both $Y = 1$ or $Y = 0$ but viewed from the conditional perspective.

To clarify this consider the likelihood conditional on $Y = a$:

$$\begin{aligned} L(\gamma, \eta \mid \mathbf{x}, \mathbf{g}, \mathbf{Y} = \mathbf{a}) &= \prod_{i=1}^{n} f_\theta(g_i \mid x_i, y_i = a) \\ &= \prod_{i=1}^{n} \frac{f_{\theta_R}(y_i = a \mid x_i, g_i)}{E_{G_{\theta_G}}(f_{\theta_R}(y_i \mid x_i, G))} f_{\theta_G}(g_i) \end{aligned}$$

For binary $Y$ we can regroup the product according to the cases: $y_i = a, g_i = b$ for $a, b \in \{0, 1\}$; we name these groups : $T_1 = \{i : y_i = 1, g_i = 1\}$ with $\|T_1\| = n_1$,$T_2 = \{i : y_i = 1, g_i = 0\}$ with $\|T_2\| = n_2$,$T_3 = \{i : y_i = 0, g_i = 1\}$with $\|T_3\| = n_3$ and finally $T_4 = \{i : y_i = 0, g_i = 0\}$with $\|T_4\| = n_4$. The consideration of these four outcome groups is important for the analyses presented below.

$$L(\gamma, \eta \mid \mathbf{x}, \mathbf{g}, \mathbf{y}) \;=\; \prod_{i \in T_1} f_\theta(1 \mid x_i, 1) \cdot \prod_{i \in T_2} f_\theta(1 \mid x_i, 0) \cdot \prod_{i \in T_3} f_\theta(0 \mid x_i, 1) \cdot \prod_{ß \in T_4} f_\theta(g = 0 \mid x_i, 0)$$

The upper limits of the sums are included to emphasize that the size of these groups are different and that conditional on the observations the size of each group are fixed. Taking logarithms where $log(f_\theta(b \mid x_i, a)) = l_\theta(b \mid x_i, a)$:

$$l(\gamma, \eta \mid \mathbf{x}, \mathbf{g}, \mathbf{y}) \;=\; \sum_{i \in T_1}^{n_1} l_\theta(1 \mid x_i, 1) + \sum_{i \in T_2}^{n_2} l_\theta(1 \mid x_i, 0) + \sum_{i \in T_3}^{n_3} l_\theta(0 \mid x_i, 1) + \sum_{i \in T_4}^{n_4} l_\theta(0 \mid x_i, 0)$$

We can expand each $l_\theta(b \mid x_i, a)$ for the \$i^{th} observation by recognizing that $f_\theta(b \mid x_i, a) = \frac{f_\theta(a \mid x_i, b)}{E_G(f_\theta(a \mid x_i, G)))}$, so that:

$$l_\theta(b \mid x_i, a) = l_\theta\left(a \mid x_i, b\right) - log\left(f(1) \cdot f_\theta(a \mid x_i, 1) + f(0) \cdot f_\theta(a \mid x_i, 0)\right)$$

Using a shorthand notation for $l_\theta(b \mid x_i, a)$ We can now input the expressions for the logistic model to obtain $\frac{\partial l_i(\theta)}{\partial \theta}$:

$$\frac{\partial l_j(\gamma, \eta)}{\partial \gamma} = \sum_{i \in T_j}^{n_j} \frac{1}{l_j(\gamma, \eta \mid x_i, g_i = b, y_i = a)} \frac{\partial l_j(\gamma, \eta)}{\partial \gamma} - \frac{\partial l_j}{\partial \gamma} log(E_G(f_{\gamma,\eta}(y_i = a \mid x_i, G)))$$

A similar expression holds for derivatives with respect to $\eta$. We present the detailed forward model expressions for the logistic model and the derivatives below:

$$f_{\theta_R}(y_i \mid x_i, g_i) = \frac{e^{\alpha + \beta x_i + \gamma g_i + \eta x_i g_i}}{1 + e^{\alpha + \beta x_i + \gamma g_i + \eta x_i g_i}}$$

To emphasize we how this works out in the logistic case:

$$l_j(\gamma, y_i = a \mid x_i, g_i = b) = \alpha + \beta x_i + \gamma b + \eta b \cdot x_i - log(1 + e^{\alpha + \beta x_i + \gamma b \eta b \cdot x_i})$$

Another convenient way to approach the problem computationally is to consider each of the four groups described above. Each group $T_j$ for the 4 possible combinations of binary $y$ and binary $g$ the likelihood contribution of each observation in each respective group has the form of :

$$f_\theta(b_j \mid x_i, a_j) = R_{j,\theta}(y = a_j, g = b_j, x_i) f_{\theta_G}(b_j)$$

Where the $i^t h$ term has the form $R_\theta(a_j, b_j, x_i) = \frac{f_{\theta_R}(a_j \mid x_i, b_j)}{E_{\theta_G}(f_{\theta_R}(a_j \mid x_i, G)}$). Because each observation contributes equally to the $f_{\theta_G}(b_j)$ this term does not impact the maximization procedure for estimation of $\gamma, \eta$. Therefore, taking logarithms of these $R_j$ and differentiating with respect to $\gamma$ and $\eta$ leads to a maximum likelihood estimation procedure. Below we present in detail the derivatives of the $R_j$ for logistic model and labeling the four groups as described above:

**Derivatives of $r_1$ for $(\gamma, \eta)$**

$$\frac{\partial r_1}{\partial \gamma} = \frac{-1 + \omega}{1 + e^{\alpha + \gamma + X(\beta + \eta)} - \omega + e^{\gamma + X\eta}\omega}$$

$$\frac{\partial r_1}{\partial \eta} = \frac{X(-1 + \omega)}{1 + e^{\alpha + \gamma + X(\beta + \eta)} - \omega + e^{\gamma + X\eta}\omega}$$

**Derivatives of $r_2$ for $(\gamma, \eta)$**

$$\frac{\partial r_2}{\partial \gamma} = -\frac{e^{\gamma+X\eta}(1+e^{\alpha+X\beta})\omega}{(1+e^{\alpha+\gamma+X(\beta+\eta)})(1+e^{\alpha+\gamma+X(\beta+\eta)}-\omega+e^{\gamma+X\eta}\omega)}$$
$$\frac{\partial r_2}{\partial \eta} = -\frac{e^{\gamma+X\eta}(1+e^{\alpha+X\beta})X\omega}{(1+e^{\alpha+\gamma+X(\beta+\eta)})(1+e^{\alpha+\gamma+X(\beta+\eta)}-\omega+e^{\gamma+X\eta}\omega)}$$

**Derivatives of $r_3$ for $(\gamma, \eta)$**

$$\frac{\partial r_3}{\partial \gamma} = \frac{e^{\alpha+\gamma+X(\beta+\eta)}(-1+\omega)}{1-e^{\alpha+\gamma+X(\beta+\eta)}(-1+\omega)+e^{\alpha+X\beta}\omega}$$
$$\frac{\partial r_3}{\partial \eta} = \frac{e^{\alpha+\gamma+X(\beta+\eta)}X(-1+\omega)}{1-e^{\alpha+\gamma+X(\beta+\eta)}(-1+\omega)+e^{\alpha+X\beta}\omega}$$

**Derivatives of $r_4$ for $(\gamma, \eta)$**

$$\frac{\partial r_4}{\partial \gamma} = -\frac{e^{\alpha+\gamma+X(\beta+\eta)}(1+e^{\alpha+X\beta})\omega}{(1+e^{\alpha+\gamma+X(\beta+\eta)})\left(-1+e^{\alpha+\gamma+X(\beta+\eta)}(-1+\omega)-e^{\alpha+X\beta}\omega\right)}$$

$$\frac{\partial r_4}{\partial \eta} = -\frac{e^{\alpha+\gamma+X(\beta+\eta)}(1+e^{\alpha+X\beta})X\omega}{(1+e^{\alpha+\gamma+X(\beta+\eta)})\left(-1+e^{\alpha+\gamma+X(\beta+\eta)}(-1+\omega)-e^{\alpha+X\beta}\omega\right)}$$

Implementation of a root finding procedure to use these derivatives to find the extrema of the log-likelihood function are provided in the GitHub repository.

### Liability Model, Binary G

The Causal Pivot is general in structre and also applicable to a continuous outcome. We consider the conditional distribution for a continuous variable when the data is obly observed conditional on being above or below a certain cutoff value. This is given by renomralizing to the tail probability of the r.v.:

$$f(y \mid I_{Y>\delta}=1, x, g) = \begin{cases} \frac{f(y|x,g)}{1-F(\delta|x,g)} & Y > \delta \\ 0 & Y \leq \delta \end{cases}$$

$$f(y \mid I_{Y>\delta}=0, x, g) = \begin{cases} \frac{f(y|x,g)}{F(\delta|x,g)} & y < \delta \\ 0 & y \geq \delta \end{cases}$$

For continuous $Y$ we can regroup the product according to the cases using an indicator variable for the event $y_i \geq \delta$ as $I_{y_i \geq \delta} = a, g_i = b$ for $a, b \in \{0, 1\}$; we name these groups : $T_1 = \{i : a_i = 1, g_i = 1\}$ with $\|T_1\| = n_1$,$T_2 = \{i : a_i = 1, g_i = 0\}$ with $\|T_2\| = n_2$,$T_3 = \{i : a_i = 0, g_i = 1\}$ with $\|T_3\| = n_3$ and finally $T_4 = \{i : a_i = 0, g_i = 0\}$with $\|T_4\| = n_4$ Then we can appluy the same Bayes rule factorization as described above and proceed with the likelihood analysis. The log likelihood has a familiar form but is more complex. To execute the analysis we again break the data into the four groups and consider the derivatives wrt $\gamma$ and $\eta$ in each group. Code executing this for a normal liability model is contained in the Github site.

#### Causal Pivot Fisher Information

Interval estimation and analysis of the limiting variance of MLE is a central advantage of a likelihood approach. We consider the Fisher Information properties of the Causal Pivot. We consider the likelihood as a random function of the stochastic system $X, G, Y$. To execute analysis we introduce a new random variable $T_{i,j}$ which takes the values $0, 1$. We consider the four cases for $Y \in A$,$Y \in A^c$,$G = 0$,$G = 1$. We let $T_1 = 1$ indicate $Y \in A, G = 1$ and 0 otherwise. $T_2 = 1$ corresponds to the state $Y \in A, G = 0$, $T_3 = 1$ corresponds to $Y \in A^c, G = 1$ and $T_4$ to $Y \in A^c, G = 0$.

$$l(\gamma, \eta \mid X, G, Y) \quad = \quad \sum_{i=1}^{n}\sum_{j=1}^{4} T_{i,j} l\left(\gamma, \eta \mid G_i = b, X_i, Y_i = a\right)$$

The Fisher Information can be computed by calculation of the quantity:

$$E\left(\begin{pmatrix} \frac{\partial l}{\partial \gamma} \\ \frac{\partial l}{\partial \eta} \end{pmatrix} \begin{pmatrix} \frac{\partial l}{\partial \gamma} \frac{\partial l}{\partial \eta} \end{pmatrix}\right)$$

This expands to this matrix:

$$I(\gamma, \eta) \quad = \quad \begin{pmatrix} E\left(\frac{\partial l}{\partial \gamma}^2\right) & E\left(\frac{\partial l}{\partial \gamma}\frac{\partial l}{\partial \eta}\right) \\ E\left(\frac{\partial l}{\partial \gamma}\frac{\partial l}{\partial \eta}\right) & E\left(\frac{\partial l}{\partial \eta}^2\right) \end{pmatrix} \tag{1}$$

If we examine a single observation from the loglikehood and consider the observations to be i.i.d, we can supress indexing in $i$:

$$l(\gamma, \eta \mid G, X, Y) = \sum_{j=1}^{4} T_j \cdot l(\gamma, \eta \mid G = b_j, X, Y = a_j)$$

Taking derivatives, generically denoting with the $'$ notation, we have:

$$l'(\gamma, \eta \mid G, X, Y) = \sum_{j=1}^{4} T_j \cdot l'(\gamma, \eta \mid G = b_j X, Y = a_j)$$

Focusing on $\gamma$, squaring this and notating $G = bj$ with $b_j$, $Y = a_j$ with $a_j$ we have:

$$\left(\frac{\partial}{\partial \gamma} l(\gamma, \eta \mid G, X, Y)\right)^2 = \left(\sum_{j=1}^{4} T_j \cdot l'_\gamma(\gamma, \eta \mid b_j \mid X, a_j)\right)^2$$

Expanding the square we see we have:

$$\left(\frac{\partial}{\partial \gamma} l(\gamma, \eta \mid G, X, Y)\right)^2 \quad = \quad \sum_{j=1}^{4} T_j \left(l'(\gamma, \eta \mid b_j \mid X, a_j)\right)^2 + 2 \cdot \sum_{j=1}^{4}\sum_{k<j} T_j T_k l'(\gamma, \eta \mid b_j, X, a_j) l'(\gamma, \eta \mid b_k X, a_k)$$

This follows because $T_j$ is an indicator variable and therefore $T_j^2 = T_j$. We also note that $T_j T_k = 0$ for $j \neq k$; therefore $E(T_j T_k h(X)) = 0$ for any $h()$. It follows that:

$$E\left(\frac{\partial}{\partial \gamma} l(\gamma, \eta \mid G, X, Y)\right)^2 \quad = \quad \sum_{j=1}^{4} E\left(T_j \left(l'_\gamma(\gamma, \eta, \mid b_j X, a_j)\right)^2\right)$$

To compute the expectation

$$E\left(T_j\left(l'_\gamma(\gamma,\eta\mid b_j,X,a_j)\right)^2\right)$$

we will condition on $G, Y$:

$$E\left(\frac{\partial}{\partial\gamma}l(\gamma,\eta\mid G,X,Y)\right)^2 \quad = \quad \sum_{j=1}^{4}E\left(E\left(T_j\left(l'_\gamma(\gamma,\eta\mid b_jX,a_j)\right)^2\mid G,Y\right)^2\right)$$

We note that the cases for T_{j} are:

$$T_j=\begin{cases}1 & G=b_j, Y=a_j\\ 0 & G\neq b_j, Y\neq a_j\end{cases}$$

So that the key quantities are:

$$E_{X\mid G=b_j,Y=a_j}\left(l'_\gamma(b_j\mid X,a_j)^2\right)$$

Explicitly:

$$E\left(\frac{\partial}{\partial\gamma}l(\gamma,\eta\mid X,G,Y)\right)^2 \quad = \quad \sum_{j=1}^{4}E_{G,Y}\left(E_{X\mid G=b_j,Y=a_j}\left(l'_\gamma(\gamma,\eta,\mid b_j,X,a_j)^2\right)\right)$$

The outer expectation is taken with respect to the joint distribution of $G, Y$. Althout both the joint distribution $f_\theta\left(G,Y\right)$ and the conditional distributions $f_\theta\left(X\mid Y,G\right)$ are calculable analytically, the observed information is known to be an excellent approximation in paractice. In this case the expectations are taken with respect to the empirical distributions of the groups:

$$E_{X\mid G=b_j,Y=a_j}\left(l'_\gamma(\gamma,\eta\mid b_j,X,a_j)^2\right)\approx\frac{1}{n_j}\sum_{T_{i,j}=1}l'\left(\gamma,\eta\mid b_j,x_i,a_j\right)^2$$

The outer expectation is taken with respect to the empirical frequency of the groups. Considering both cases and controls this is $p_j=\frac{n_j}{n}$. In the instance of cases only inference this is $p_j=\frac{n_j}{n_c}$.

The overall Fisher information of a sample is $n_sI(\theta)$ where $n$ is representing the sample size; $s$ is used to indicate that this can be considered in the cases only of case-control context. Multiplying by $n_s$ we are left with :

$$I_s(\theta)=\sum n_jI_j\theta(a)$$

Confidence intervals are formed by computing the asymptotic variance of the MLE as $I^{-1}(\theta) = \left(\sum_j n_jI_j\left(\theta\right)\right)^{-1}$. Let $\sigma^2_\gamma$ be the (1,1) entry in the matrix $I^{-1}(\theta)$ for a given sample. The interval boundaries for $\gamma$ are determined $P(Z_{\frac{\alpha}{2}}\leq\frac{\hat{\gamma}-\gamma}{\sigma_\gamma}\leq Z_{1-\frac{\alpha}{2}})=1-\alpha$ so that the intervals are $\hat{\gamma}\pm\sigma_\gamma\cdot Z_{\frac{\alpha}{2}}$.
