## SupplementaryAnalyses for "The Causal Pivot: A Structural Approach to Genetic Heterogeneity and Variant Discovery in Complex Diseases"

**Supplementary Results**

**Overview**

This Supplement presents additional results related to the Causal Pivot. The first section contains results for the impact of PRS effect size on conditional means in the case of binary trait outcomes. Next, details are presented concerning power analyses for the CP-LRT in the continuous trait liability. We then provide additional details related to the analysis of UKB data. These include characterization of the PRS used in analyses and enumeration of sequence ontology for the variants that were used to designate the status of RV+ ; variants are enumerated according to variant types for the demonstration disease genes. Negative control analyses are also presented using rare synonymous variants in the demonstration genes. We also provide more detailed analyses of CP-LRT performance using permutation of RV+ to determine the contours of the CP-LRT MLE estimates in UKB data under the null hypothesis of no RV effect. We then present results for the inverse probability weighting method to correct for ancestry confounding in UKB data using random forest models.

**The Impact of PRS Effect Size**

We studied the impact of the PRS effect size on the PRS-RV causal pivot. We focus on the case of a binary phenotype using the logistic outcome model. We treat the PRS as normally distributed and the RV as a Bernoulli trial. Using the mathematical derivation of conditional distributions presented in the Methods and as described further in the Mathematical Supplement, the conditional means of both the PRS and RV can be determined. Subsequently, the conditional means can be considered as a function of the strength of the PRS effect on trait outcome. The result for the conditional mean E(G|X,Y) is presented in terms of conditional odds ratio that G=1 – that is the chance for an individual to bear a functional RV.

**Supplemental Figure 1**

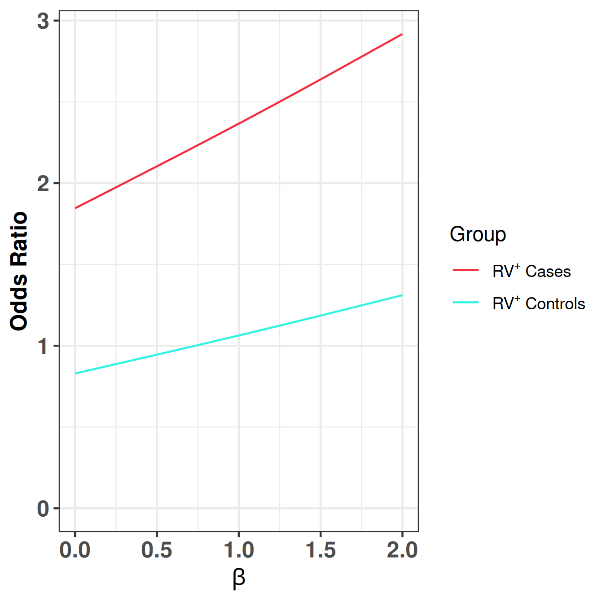

**Suppl. Fig 1: Odds Ratio of RV+ as a function of PRS effect size: the dynamics of RV risk.** The figure presents the effect of variation in PRS effect size (x-axis), denoted β. The log-odds transformation is used to determine an odds ratio of RV+ comparing the situation of PRS values of X=1 and X=-1; this odds ratio value is indicated by the Y-axis of the plot. The odds are defined: E(G |X,Y)/(1-E(G|X,Y) . The additional model parameters are fixed: α=−2.2; γ=1; η=−.4. The red line presents the case of disease where Y=1 (Cases), and the blue line presents controls where Y=0. For cases (Y=1), the odds ratio RV+ when X=-1 vs X=1 is greater than 1 and increasing in β. This shows that as the explanatory power of the PRS increases the relative likelihood of RV+ when the PRS is low vs when the PRS is high becomes increasingly elevated. The result among controls is more modest and the OR remains close to 1 as a function of β. This indicates that among controls the impact on the odds of RV+ is modestly dependent on PRS, but there is an effect of PRS effect size.

We also examined the impact of the PRS effect on the conditional mean of the PRS, denoted X, in the disease and control groups under the logistic model. There are four conditional mean functions corresponding to the four states: RV+,Y=1; RV-,Y=1; RV+,Y=0,RV-,Y=0.

**Supplemental Figure 2**

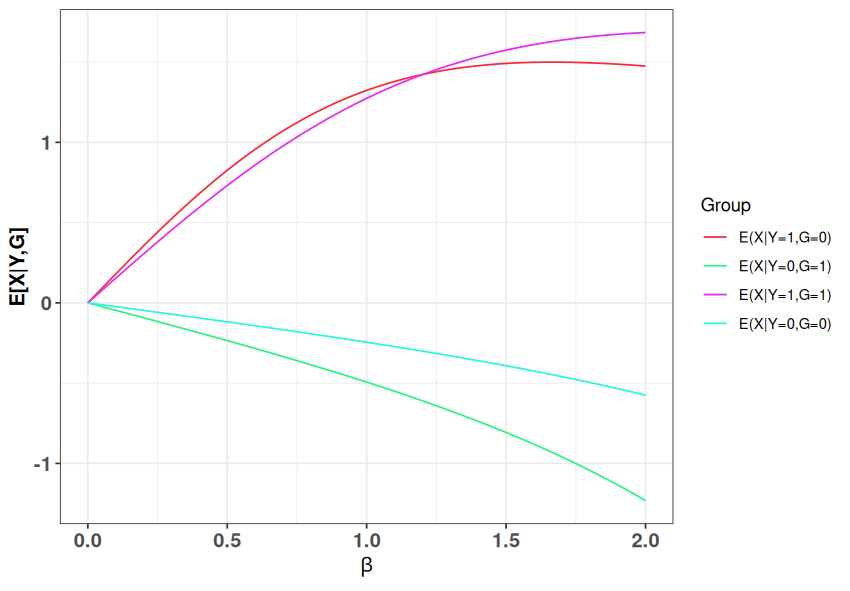

**Suppl. Fig 2: Conditional mean of the PRS as a function of PRS effect size: asymmetry between cases and controls.** The figure presents the asymmetric relationship of E(X |G,Y) across the four conditional states of G and Y. Each curve presents the conditional mean (y-axis) as a function of PRS effect size β (x-axis). The other parameters of the forward outcome model are fixed: α=−2.2; γ=1; η=−.4. The conditional means are presented as four distinct curves. Interestingly, when Y=1, the conditional means are similar functions of β ; they diverge modestly as β increases, but they cross at very high values – in this case around β=1.25. This shows that among cases the conditional mean of the PRS increases as a function of PRS effect size, but the pattern is highly similar among the RV+ and RV- cases. For controls the pattern is a decreasing function of β. Interestingly, there is a divergence in the conditional mean of the PRS given RV+ and RV- among the controls (Y=0) that increases as β becomes stronger.

**Liability Model for Quantitative Y**

Power analyses were performed by simulating observations from the linear trait model and then selecting as observable the values over a set quantile cutoff. We used a quantile of 0.92 driven by the UKB data for an LDL cholesterol level over 190. CP-LRT analyses were performed using the likelihood ratio test procedure as described in the main text for the liability model. Maximum likelihood estimates are determined by score equations using the conditional trait density function.

Traits were simulated using a linear model with α = 0, β = 0.3, and RV frequency ω = 0.001. Conditional outcomes were created by selecting individuals exceeding the 93rd percentile of the trait distribution, (LDL direct > 4.9 nmol/L) as an example clinical hypercholesterolemia threshold. Comparisons include CP-LRT, Wilcoxon test, Z-test, and unconditional linear regression using the full sample. CP-LRT showed superior power over the Wilcoxon test in the absence of interaction effects, while the Z-test and linear regression achieved higher power overall. Interaction effects increased power across all methods, with the Wilcoxon test showing the largest improvement.

**Supplemental Figure 3**

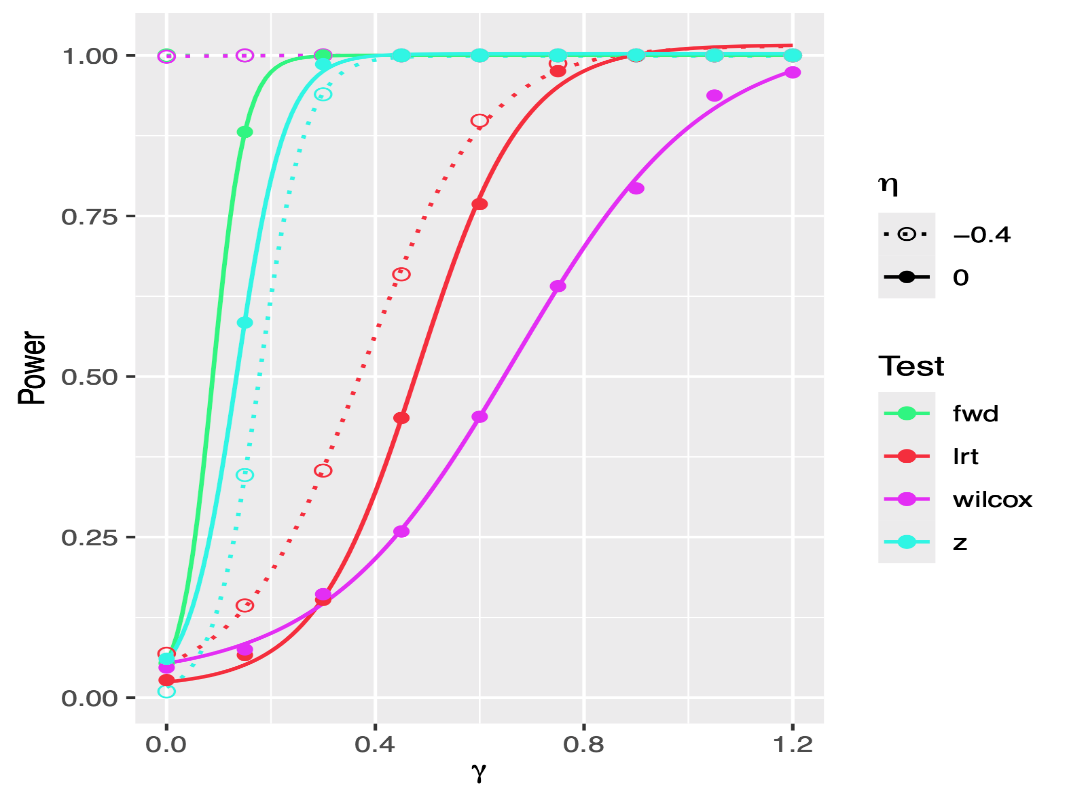

**Suppl. Fig 3 Liability Model Power Analyses: a comparison of methods** The figure presents CP-LRT power analyses for the liability model. The x-axis is the strength of the RV effect – parameterized as γ.The values of the other parameters are held constant: ω = 0.001, α = 0, β = 0.4, and trait cutoff δ = 1.33 corresponding to the trait quantile 0.92. We set μ_X_= 0, σ_X_= 1. The figure shows that CP-LRT power is strongly higher than the Wilcoxon in the absence of a PRS-RV interaction; however the Z-test has better power than the CP-LRT. The forward regression using all of the data has the highest power. When an interaction effect is present the power is elevated for all tests; the effect is most pronounced for the Wilcoxon whose power shifts from below the CP-LRT to above.

**UKB Analyses**

**Polygenic Risk Scores**

Polygenic scores were extracted for the subjects using the available data in the UKB RAP. These scores are normal in the disease population (Suppl. Figure 4).

**Supplementary Figure 4**

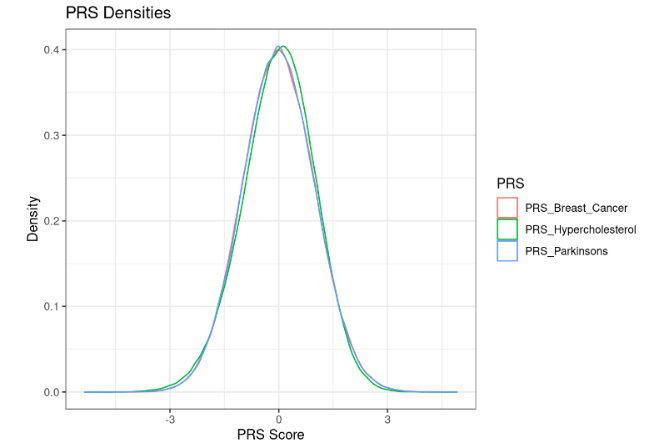

**Supp Fig.4: Evidence of normality of UKB PRS: an expected result.** PRS score distributions from the sample metadata after removal of samples for which we do not have all PRS scores. Additionally any samples that had more than one of the phenotypes were excluded from the target set (i.e BC+ and PD+ individuals are rare but excluded). The population distribution of scores are well approximated by Gaussian distributions. We shifted and scaled these distributions to have mean 0 and variance 1.

**Classify Pathogenic Variants**

Variants were selected to determine the RV+ status of individuals. Variants were considered positive if they were classified as Clinvar Pathogenic OR a rare (MAF <0.01) stop-gain or rare frameshift. The rare status was defined by gnomAD v3 non-Finnish European allele frequencies with frequency less than 0.01. Clinvar Pathogenic was defined as either explicitly Pathogenic or Likely_pathogenic OR conflicting_interpretations_of_pathogenicity with at least one pathogenic or likely pathogenic and no benign or likely benign submission. We further excluded pathogenic markers with qualifiers such as “drug_response” or “risk_factor”.

**Exclude positive markers strongly correlated w/ PRS**

To ensure independence of PRS and rare variants we used logistic regression to find positive events in genes of interest that are significantly correlated with their associated PRS. Such variants were flagged for exclusion from pathogenic classification. Included variants are shown in Supplemental File 3.

**Supplementary Figure 5**

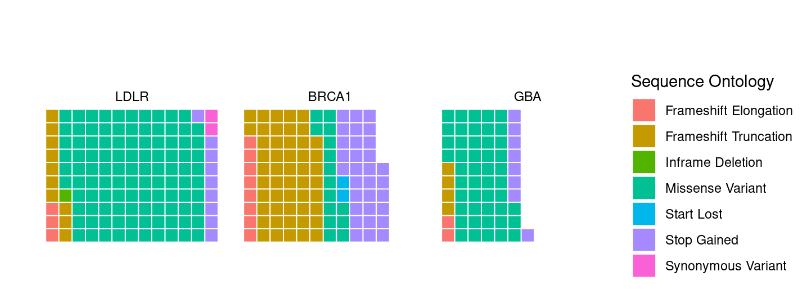

**Suppl Fig.5. Variant mutation type waffle plot: diversity of variants for each disease gene.** The plot shows the visual representation of different variants by type for each disease gene. The diversity and number of variants emphasizes that the PRS conditional effects are not driven by a single variant type.

**MLE Analysis**

We used the CP Maximum Likelihood Estimation (MLE) procedure to estimate γ (the coefficient for G) and η (the coefficient for X * G) of the reverse logistic model then calculate the chi-squared test statistic for the resulting maximum Log Likelihood (LL). We solved for the maximum LL via optim() using the build-in SANN simulated annealing algorithm. To correct for variability in some areas of the parameter space, we solved the optimization multiple times, then took the median result for the parameter estimate.

**Supplementary Table 1**

| Cohort | Gene | α | β | ω |
| --- | --- | --- | --- | --- |
| HC190 | *LDLR* | -2.86 | 0.723 | 0.00216 |
|  | *BRCA1* | -2.86 | 0.723 | 0.00088 |
|  | *GBA* | -2.86 | 0.723 | 0.00237 |
| BC | *LDLR* | -2.93 | 0.626 | 0.00236 |
|  | *BRCA1* | -2.93 | 0.626 | 0.000769 |
|  | *GBA* | -2.93 | 0.626 | 0.0022 |
| PD | *LDLR* | -4.96 | 0.404 | 0.00217 |
|  | *BRCA1* | -4.96 | 0.404 | 0.000882 |
|  | *GBA* | -4.96 | 0.404 | 0.0024 |

**Suppl Table 1 Parameters supplied for CP-LRT Analyses.**  Nine CP-LRT were performed; three tests for each disease gene RV status across the three disease PRS cohorts. Cohort represents the UKB phenotype and PRS selection, Gene identifies the RVs. All parameters are derived from analysis of UKB data restricting to individuals with exome data from European ancestry. BC analyses are restricted to females. The α parameter is the disease prevalence; β is the PRS effect; ω is the RV frequency.

**Supplementary Figure 6**

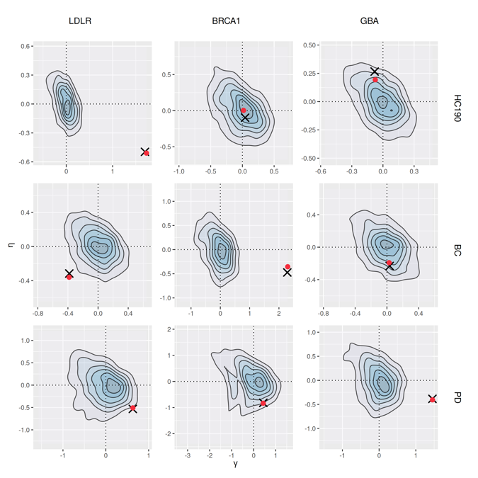

**Suppl Figure 6: Permutation contour plot of the CP-LRT procedure: concordance with the forward regression results** The graph shows MLE results for the CP-LRT for γ on the x-axis and η on the y-axis. The contour are the results of random permutation analyses where the RV status is randomized in the respective disease cohort; results center at or near the point (0,0), as expected; there is a negative association of the γ, η parameter estimates, as shown under the Fisher Information analysis. The red dots indicate the results obtained in cases only analyses; the **x** symbols identify the parameter estimate from fitting a full case-control forward logistic regression model for disease outcome as binary with the PRS and RV variables as explanatory factors where η is their interaction effect. The cases-only CP MLE is similar to the case control result as evidenced by the overlap of the red dot with the **x.**

**Supplementary Table 2. CP-LRT logitG Model Parameter Estimates with Confidence Intervals**

| **Cohort** | **Gene RV+** |  |  | **95% CI** |  |  | |  | | **95% CI** | |
| --- | --- | --- | --- | --- | --- | --- | --- | --- | --- | --- | --- |
|  |  | CPLRT P-Value | Estimate γ | LL | UL |  | Estimate η | | LL | | UL |
| Hypercholesterolemia | *LDLR* | 0 | 1.729464 | 1.538257 | 1.920672 |  | -0.50931 | | -0.72972 | | -0.2889 |
|  | *BRCA1* | 0.859 | 0.017978 | -0.55608 | 0.592033 |  | 0.003322 | | -0.64334 | | 0.649982 |
|  | *GBA* | 0.453 | -0.07402 | -0.41859 | 0.270559 |  | 0.192979 | | -0.13378 | | 0.519733 |
| Breast Cancer | *LDLR* | 0.024 | -0.38346 | -0.87308 | 0.106153 |  | -0.35973 | | -0.74722 | | 0.027762 |
|  | *BRCA1* | 0 | 2.298798 | 1.891172 | 2.706425 |  | -0.35441 | | -0.84491 | | 0.136081 |
|  | *GBA* | 0.59 | 0.027694` | -0.41155 | 0.466939 |  | -0.19154 | | -0.62661 | | 0.243526 |
| Parkinson's Disease | *LDLR* | 0.058 | 0.631394 | 0.028685 | 1.234104 |  | -0.50932 | | -1.10341 | | 0.084774 |
|  | *BRCA1* | 0.215 | 0.433668 | -0.63972 | 1.507054 |  | -0.835 | | -1.85542 | | 0.185414 |
|  | *GBA* | 0 | 1.454348 | 1.063813 | 1.844884 |  | -0.40495 | | -0.84035 | | 0.030444 |

**Suppl Table 2 CP-LRT Results and Confidence Intervals** The table presents CP-LRT p-values, parameter estimates and 95% Confidence Intervals for each of the 9 anlyses performed on UKB data. The p-value is the CP-LRT result. Estimates are MLEs for γ and η; and confidence intervals are using the 0.025 and 0.975 quantiles of the normal distribution multiplied against the square root of the diagonal elements of the asymptotic covariance matrix of the MLEs as estimated from the observed Fisher information derived from the CP maximum likelihood analysis.

**The Specificity of the CP Effect Depends on Pathogenic Variants**

We also performed analysis on rare synonymous variants using the CP-LRT procedure we applied to rare pathogenic or suspected pathogenic variants. We applied our same tertile procedure to these variants, and the results are show in this figure:

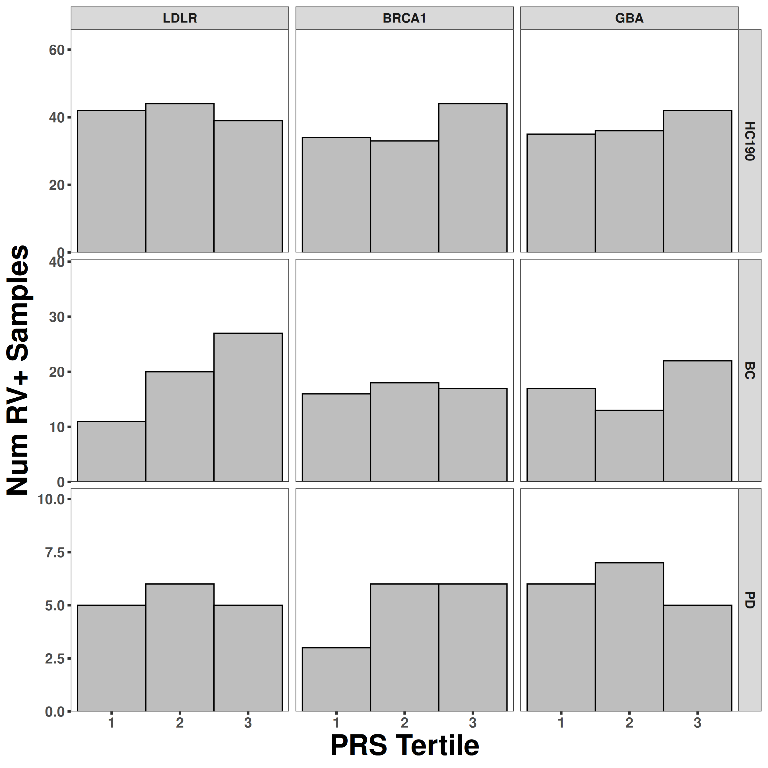

**Supplementary Figure 7: Negative control variants tertile plot.** The figure presents the tertile analysis as presented in Fig3 of the main paper but now applied to a set of synonymous rare variants. These variants do not show the negative correlation pattern observed along the diagonal in Figure 3. There are no significant CP-LRT associations detected.

**Supplementary Figure 8**

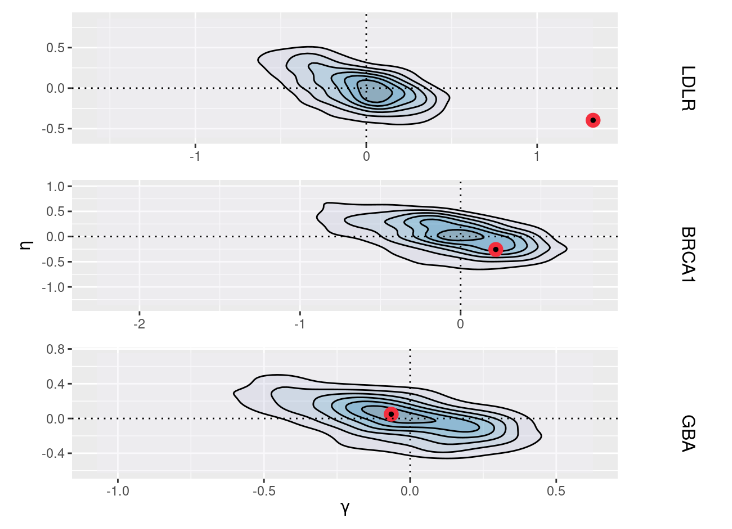

**Suppl Figure 8: Liability model contour analysis on UKB data.** The graphic shows MLE results for the CP-LRT in the liability model for the LDLR cholesterol treated as a quantitative trait using subjects with a LDL value above the 92%-ile of the trait value. Results for γ on the x-axis and η on the y-axis. The contour graphics are the results of random permutation analyses where the RV status is randomized in the disease cohort; results center at or near the point (0,0), as expected. The red dots indicate the results obtained in cases only analyses. Analyses were performed using LDLR variants as well as BRCA1 and GBA1l the BRCA1 and GBA1 analyses serve as negative controls.

**Ancestry Balancing via Inverse Probability Weighting**

Here, we perform an alternative method of ancestry balancing using inverse probability weighting. Propensities are calculated using random forests over the first 15 PCs from UKB. Again, we calculate a permutation distribution; our statistic is the difference in weighted PRS sums between RV+ and RV- cases. We also present feature “importance” metrics.

**Supplementary Figure 9**

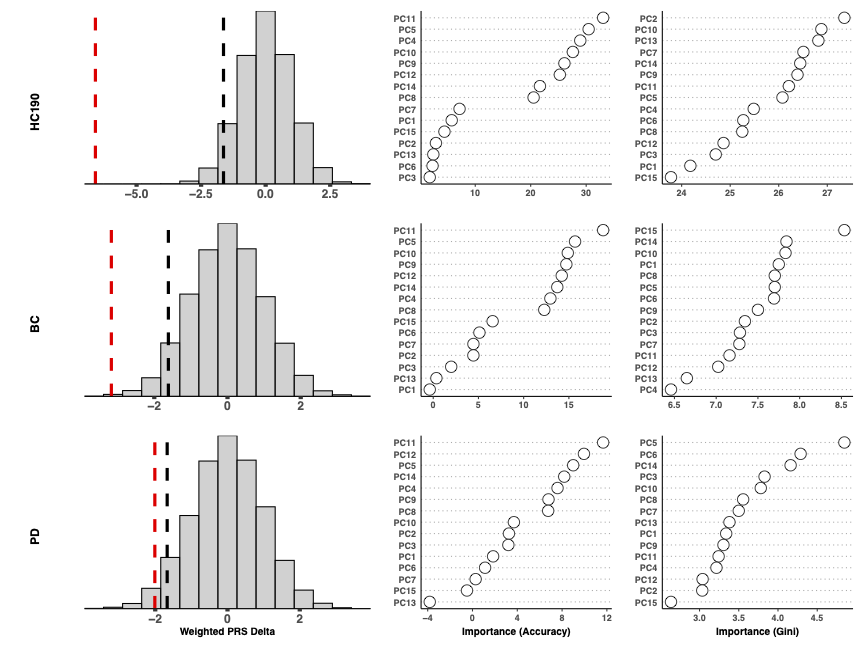

**Suppl Figure 9: Inverse probability weighting to control for ancestry: a machine learning approach.** We employed random forest models to determine a propensity function for RV+ status among cases for each disease and its respective disease gene. Each row represents a distinct disease cohort. The left column of plots shows the result for the IPW difference of RV+ and RV- cases by the red dashed vertical line; a negative value indicates that the IP weighted difference between RV+ and RV- cases. The histogram shows the results of permutation assignment of RV+ status in each cohort; the black dashed vertical line is the 0.05 quantile of the permutation distribution. The second and third columns in each row present the random forest importance plots that depict the relevance of the respective ancestry principal components in the random forest prediction. Interestingly, high order components are predictive, indicating that the rare RV occur in sublineages of the population tagged by ancestry components that explain less global ancestry. This observation is consistent with the pattern that RV+ status is distinct from the common variation that drives ancestry PCA.
