## Supplementary material for "The Causal Pivot: A Structural Approach to Genetic Heterogeneity and Variant Discovery in Complex Diseases": CodeSupplement: Code_Supplement_20241214.html

Causal Pivot Analysis


### Causal Pivot Analysis

###### 13 December, 2024, 16:35

- 1 Causal Pivot Analysis -
  Introduction
- 2 Setup
  - 2.1 Initialize Data and
    Constants
  - 2.2 Load Dependencies
  - 2.3 Load PRS
    - 2.3.1 Remove samples w/o PRS from
      cohort.meta
  - 2.4 Define Cohorts
  - 2.5 Classify Pathogenic
    Variants
  - 2.6 Flag markers strongly correlated
    w/ PRS
  - 2.7 Create genomic event matrices
    - 2.7.1 Positive (rare pathogenic)
      markers
    - 2.7.2 Control (rare synonymous)
      markers
  - 2.8 Create master dataframe &
    cohort data frames
    - 2.8.1 Pathogenic Variants
    - 2.8.2 Control Variants
- 3 Rare Variant Summaries
  - 3.1 Create publishable variant Excel
    workbook
  - 3.2 Plot variant classification
    summaries
    - 3.2.1 Variant mutation type waffle
      plot
    - 3.2.2 Variant spread hisogram
- 4 Perform MLE Permutation Testing
  - 4.1 Set Permutation Count
  - 4.2 Logit G
  - 4.3 Liability G
- 5 Plot Collider Effect
  - 5.1 Plot Case + Control Scatter
    Plot
    - 5.1.1 Pathogenic Variants
  - 5.2 Plot collider effect as PRS
    tertiles bar plot
    - 5.2.1 Pathogenic Variants
- 6 Compose UKBB Collider Summary
  Plot
- 7 Perform Ancestry Confounder Control
  Analysis
  - 7.1 Ancestry k Nearest Neighbors
    analysis
  - 7.2 Ancestry Balancing via Inverse
    Probability Weighting
- 8 Supplementary LSD/PD “Genetic Load”
  Analysis

### 1 Causal Pivot Analysis - Introduction

This document contains code and comments for the primary analysis of
the presence of a causal pivot between two independent causal factors of
disease:

- Pathogenic Rare Variant Events (RV) - a binary variable
- Polygenic Risk Scores (PRS) - a continuous variable

3 disease cohorts from the UK Biobank (UKB), and associated causative
genes are defined:

- Hypercholesterolemia & LDLR
- Breast Cancer & BRCA1
- Parkinson’s & GBA

Each cohort/gene is paired with the “standard” version of the UKB PRS
for that disease.

Running the code step-by-step and reading the comments should be the
simplest way to understand the analysis.

The analysis proceeds as follows:

1. partition UKB samples into sub-cohorts based on disease
   metadata
2. classify rare/pathogenic variants
3. adjust for population-wide dependence b/w rare variant status and
   PRS
4. create binary rare event variables
5. investigate causal collider via plots and statistics
6. investigate presence of collider signal after controlling for
   ancestry
7. perform supplemental “genetic load” analysis in Parkinson’s

- the Lysosomal Storage Disease (LSD) gene list is required and read
  from `LSD_human_genes.txt` in `SRC_DATA_DIR`.

### 2 Setup

#### 2.1 Initialize Data and Constants

**NOTE:** this can take a while…

Initialize analytic products from variant extraction workflow. See init.R for initial data transforms. This creates
multiple data objects and saves them in the configured
`SRC_DATA_DIR`:

- `cohort.meta.rds`: metadata for European UKB samples from
  the RAP cohort browser
- `annotations.rds`: variant annotations from
  open-cravat
- `geno.mtx.rds`: genotypes as a sparse
  `dgCMatrix` matrix

The analysis will re-use pre-existing exist files if they exist. To
run an analysis from a new set of objects, set a new
`SRC_DATA_DIR` or delete the files.

#### 2.2 Load Dependencies

#### 2.3 Load PRS

Load PRS scores. Samples missing PRS data are excluded.

```
# load and format prs data ####

prs.name.map <- list(
  "Standard PRS for low density lipoprotein cholesterol (LDL_SF)" = "PRS_Hypercholesterol"
  ,"Standard PRS for breast cancer (BC)" = "PRS_Breast_Cancer"
  ,"Standard PRS for parkinson's disease (PD)" = "PRS_Parkinsons"
)

prs <- cohort.meta %>%
  filter(!if_any(all_of(names(prs.name.map)), is.na)) %>%
  select(
    sample.id,
    all_of(names(prs.name.map))
  ) %>%
  rename_with(~ unlist(prs.name.map), all_of(names(prs.name.map)))

# normalize prs distributions to mean = 0 and sd = 1 ####

prs <- prs %>%
  mutate(across(starts_with('PRS'), ~ (. - mean(.)) / sd(.)))

cohort.meta <- cohort.meta[na.omit(match(prs$sample.id, cohort.meta$sample.id)),]

# save `prs.rds` ####

saveRDS(prs, file = file.path(OUT_DIR, 'prs.rds'))

# display and save PRS densities plot ####

(
  prs %>%
  mutate(across(starts_with('PRS'), ~ (. - mean(.)) / sd(.))) %>%
  pivot_longer(cols = starts_with('PRS'), names_to = 'PRS', values_to = 'score') %>%
  ggplot(aes(x = score, col = PRS)) +
  geom_density() +
  labs(title = 'PRS Densities', x = 'PRS Score', y = 'Density') +
  theme_bw()
) %>%
  save.plot('prs.densities', root.dir = PLOTS_DIR, sub.dir = 'supplementary')
```

##### 2.3.1 Remove samples w/o PRS from cohort.meta

#### 2.4 Define Cohorts

Define relationships between cohorts, genes and PRS scores and helper
objects for later data manipulation.

Disease cohorts contain:

- gene(s) of interest
- name of column in `prs` data-frame for corresponding
  disease PRS
- case/control cohort inclusion function
- cohort exclusion function

```
# define global COHORTS list ####

COHORTS <- list(
  "HC190" = list(
    "gene.grp" = 'LDLR',
    "genes" = c('LDLR'),
    "prs" = "PRS_Hypercholesterol",
    "in.cohort" = function(cohort.meta) {
      high.ldl <- cohort.meta$ldl.inst.0 > 4.9
      !is.na(high.ldl) & high.ldl
    },
    "exclude" = function(cohort.meta) {
      is.na(cohort.meta$ldl.inst.0)
    }
  ),
  "BC" = list(
    "gene.grp" = 'BRCA1',
    "genes" = c('BRCA1'),
    "prs" = "PRS_Breast_Cancer",
    "in.cohort" = function(cohort.meta) {
      bc.inst.0 <- cohort.meta$bc.inst.0
      bc.inst.1 <- cohort.meta$bc.inst.1
      ((!is.na(bc.inst.0) & bc.inst.0) |  (!is.na(bc.inst.1) & bc.inst.1))
    },
    "exclude" = function(cohort.meta) {
      cohort.meta$sex != 'F'
    }
  ),
  "PD" = list(
    "gene.grp" = 'GBA',
    "genes" = c('GBA'),
    "prs" = "PRS_Parkinsons",
    "in.cohort" = function(cohort.meta) {
      !is.na(cohort.meta$pd.rep.date)
    }
  )
) %>%
lapply(function(cohort) {
  if (!is.null(cohort$exclude)) {
    cohort.meta <- cohort.meta[!cohort$exclude(cohort.meta),]
  }
  cohort$labels <- set_names(cohort$in.cohort(cohort.meta), cohort.meta$sample.id)
  cohort
})

# remove samples in multiple cohorts ####

# create dataframe flagging samples for inclusion in our separate cohorts
cohort.flags <- purrr::map(COHORTS, ~.$labels[cohort.meta$sample.id]) %>%
  as.data.frame() %>%
  # mutate(across(everything(), ~ifelse(is.na(.), F, .))) %>%
  filter(rowSums(across(everything()), na.rm = T) < 2)

# update cohort labels to exclude multi-cohort samples
for (cohort.name in names(COHORTS)) {
  cohort <- COHORTS[[cohort.name]]
  cohort$labels <- cohort$labels[which(names(cohort$labels) %in% rownames(cohort.flags))]
  COHORTS[[cohort.name]] <- cohort
}

# Create some helper objects ####
# map cohort names to gene group labels
cohort.prs.names <- unlist(map(COHORTS, ~.$prs))

# map cohort names to gene group labels
cohort.gene.grps <- unlist(map(COHORTS, ~.$gene.grp))

# map gene groups to vector of gene labels
gene.grps <- set_names(map(COHORTS, ~.$genes), cohort.gene.grps)

# map gene label to its gene group
gene.grp.lkp <- lapply(COHORTS, function(cohort) {
  map(cohort$genes, function(x) cohort$gene.grp) %>% setNames(cohort$genes)
}) %>%
  setNames(NULL) %>%
  unlist()

# vector mapping sample name to cohort
cohort.marks <- names(COHORTS) %>%
  sapply(function(cohort) {
    rownames(cohort.flags[which(cohort.flags[[cohort]]),]) %>%
      setNames(rep_along(., cohort), .)
  }) %>%
  unname %>%
  unlist

# print cohort counts ####
cat('Num. Samples per Cohort:\n')
```

```
## Num. Samples per Cohort:
```

```
table(cohort.marks)
```

```
## cohort.marks
##    BC HC190    PD 
## 12479 24656  2940
```

#### 2.5 Classify Pathogenic Variants

Classify pathogenic markers via conservative heuristics applied to
ClinVar, gnomAD and sequence ontology annotations.

A variant is “pathogenic” if it is classified as Clinvar Pathogenic
OR a rare stopgain OR rare frameshift event.

“Rare” defined by gnomAD v3 non-Finnish European allele frequency lt.
cutoff.

A Clinvar Pathogenic annotation is defined as containing:

- a “Pathogenic” significance `Pathogenic`,
  `Likely_pathogenic`,
  `Pathogenic/Likely_pathogenic`, etc.
- AND
- NOT a `Conflicting_interpretations_of_pathogenicity`
- AND
- NOT a “qualified” significance
  (e.g. `Pathogenic|drug_response`)

Examples:

- `Pathogenic/Likely_pathogenic` => pathogenic
- `Conflicting_interpretations_of_pathogenicity` => NOT
  pathogenic
- `Pathogenic/Likely_pathogenic|risk_factor` => NOT
  pathogenic

See variant.classifiers.R
for filter functions.

```
# flag pathogenic variants in genes of interest ####

# define "rare' variant cutoff - 0.1% allele frequency
gnomad.cutoff <- 0.001

# run heuristics
rv.flags <- annotations %>%
  filter(gene %in% unlist(gene.grps)) %>%
  classify.clinvar() %>%
  classify.lof() %>%
  classify.synonymous() %>%
  classify.missense() %>%
  mutate(
    class.rare = gnomad3.af_nfe < gnomad.cutoff,
  ) %>%
  class.na.false() %>%
  mutate(
    class.positive = class.clinvar.pathogenic | (class.rare & class.loss.of.function),
    class.rare.synonymous = class.rare & class.synonymous,
    class.rare.missense = class.rare & class.missense,
  ) %>%
  select(
    marker.id,
    gene,
    class.positive,
    class.rare.synonymous,
    class.rare.missense,
  )

# save result ####

saveRDS(rv.flags, file = file.path(OUT_DIR, 'rv.flags.rds'))
```

#### 2.6 Flag markers strongly correlated w/ PRS

Show that the causal factors are independent.

Flag markers for exclusion using logistic regression to identify
markers that significantly correlated with associated PRS.

Previously processed variants are cached to improve
iterative/cumulative analysis. Delete or modify cached object to
re-process variants.

```
# read from cached RDS file if it exists ####

MARKER_PRS_CORS_RDS <- file.path(DATA_DIR, "marker.prs.cors.df.rds")

if (file.exists(MARKER_PRS_CORS_RDS)) {
  marker.prs.cors <- readRDS(MARKER_PRS_CORS_RDS)
} else {
  marker.prs.cors <- list()
}

# calculate marker/PRS associations based on cohort gene/PRS combos ####

for (cohort.name in names(COHORTS)) {
  cohort <- COHORTS[[cohort.name]]
  
  if (cohort.name == 'PD') {
    # we want to associate the LSD pathway genes with Parkinsons PRS
    # for genetic burden analysis
    cohort.marker.ids <-
      rv.flags[rv.flags$gene %in% c(cohort$genes, lsd.genes),]$marker.id
  } else {
    cohort.marker.ids <- rv.flags[rv.flags$gene %in% cohort$genes,]$marker.id
  }
  
  # filter positive variant IDs in genes of interest that we haven't processed
  if (!is.null(marker.prs.cors[[cohort.name]])) {
    cohort.marker.ids <-
      setdiff(cohort.marker.ids, marker.prs.cors[[cohort.name]]$marker.id)
  }
  
  # create logical vector for training
  cohort.marker.events <- geno.mtx[, cohort.marker.ids] > 0
  
  # apply over marker.ids to calculate correlation significance
  prs.cors <- lapply(cohort.marker.ids, function(marker.id) {
    events <- cohort.marker.events[match(names(cohort$labels), rownames(cohort.marker.events)),
                                   marker.id]
    prs.vals <-
      prs[match(names(cohort$labels), prs$sample.id), ][[cohort$prs]]
    
    data <- data.frame(event = events,
                       prs = prs.vals)
    
    my.fit <-
      suppressWarnings(glm(
        event ~ prs,
        family = binomial(link = 'logit'),
        data = data
      ))
    fit.coefs <- coef(summary(my.fit))
    c('marker.id' = marker.id,
      'cor' = fit.coefs[2, ][['Estimate']],
      'p.val' = fit.coefs[2, ][['Pr(>|z|)']])
  }) %>%
    do.call(rbind, .) %>%
    as_tibble()
  
  marker.prs.cors[[cohort.name]] <-
    rbind(marker.prs.cors[[cohort.name]], prs.cors)
}

# cache result ####

saveRDS(marker.prs.cors, MARKER_PRS_CORS_RDS)

# flag markers with significant association w/ corresponding PRS ####

prs.sig.marker.ids <- concat.df.list(marker.prs.cors, key.cols = 'cohort') %>%
  mutate(p.val = as.numeric(format(p.val, scientific = F))) %>%
  filter(p.val < 0.05) %>%
  pull(marker.id) %>%
  unique()

# print results ####

cat("# Pathogenic Markers w/ Sig. PRS. Assoc. per Gene\n")
```

```
## # Pathogenic Markers w/ Sig. PRS. Assoc. per Gene
```

```
table(gene.grp.lkp[rv.flags[rv.flags$marker.id %in% prs.sig.marker.ids & rv.flags$class.positive,]$gene])
```

```
## 
## BRCA1   GBA  LDLR 
##     4     4     9
```

```
cat("# Control Markers w/ Sig. PRS. Assoc. per Gene\n")
```

```
## # Control Markers w/ Sig. PRS. Assoc. per Gene
```

```
table(gene.grp.lkp[rv.flags[rv.flags$marker.id %in% prs.sig.marker.ids & rv.flags$class.rare.synonymous,]$gene])
```

```
## 
## BRCA1   GBA  LDLR 
##     1     3    12
```

#### 2.7 Create genomic event matrices

##### 2.7.1 Positive (rare pathogenic) markers

Create Rare Variant (`RV`) “event” matrix counting
pathogenic variants per gene.

```
# flag pathogenic markers - class.positive & !prs.associated ####

rv.flags$pathogenic <- rv.flags$class.positive &
  !(rv.flags$marker.id %in% prs.sig.marker.ids)

# create named vector of gene groups - labels are marker.ids ####

rv.gene.marks <- rv.flags %>%
  filter(pathogenic) %>%
  select(marker.id, gene) %>%
  rowwise() %>%
  mutate(gene.grp = gene.grp.lkp[[gene]]) %>%
  ungroup() %>%
  with(setNames(gene.grp, marker.id))
  

# subset geno.mtx w/ only rare pathogenic events ####

rv.mtx <- geno.mtx[,match(names(rv.gene.marks), colnames(geno.mtx))]

# count pathogenic rare variant events per gene, per sample ####

gene.rv.events <- alt.sums.by.var(rv.mtx, rv.gene.marks)

# print RV counts per gene group ####

cat("Gene Group Pathogenic RV Counts\n")
```

```
## Gene Group Pathogenic RV Counts
```

```
table(rv.gene.marks)
```

```
## rv.gene.marks
## BRCA1   GBA  LDLR 
##   106    61   130
```

##### 2.7.2 Control (rare synonymous) markers

Create Control Variant (`CV`) event matrix.

```
# flag control markers - class.rare.synonymous & !prs.associated ####

rv.flags$control <- rv.flags$class.rare.synonymous &
  !(rv.flags$marker.id %in% prs.sig.marker.ids)

# create named vector of gene groups - labels are marker.ids ####

cv.gene.marks <- rv.flags %>%
  filter(control) %>%
  select(marker.id, gene) %>%
  rowwise() %>%
  mutate(gene.grp = gene.grp.lkp[[gene]]) %>%
  ungroup() %>%
  with(setNames(gene.grp, marker.id))
  

# subset geno.mtx w/ only rare pathogenic events ####

cv.mtx <- geno.mtx[,match(names(cv.gene.marks), colnames(geno.mtx))]

# count pathogenic rare variant events per gene, per sample ####

gene.cv.events <- alt.sums.by.var(cv.mtx, cv.gene.marks)

# print RV counts per gene group ####

cat("Gene Group Control RV Counts\n")
```

```
## Gene Group Control RV Counts
```

```
table(cv.gene.marks)
```

```
## cv.gene.marks
## BRCA1   GBA  LDLR 
##    73    39    81
```

#### 2.8 Create master dataframe & cohort data frames

Collate variables into “master” dataframe and a cohort-specific list
of dataframes. These two objects are our primary analysis inputs.

The master dataframe contains one row per sample for ALL samples and
include:

- cohort inclusion flags for all three cohorts
- gene rare variant flags for all three genes
- prs values for all three cohorts

The dataframes in the list contain ONLY samples in each cohort and
include:

- `sample.id`
- `case`: disease flag (case/control flag) - forward model
  response variable
- `rv`: pathogenic rare variant event flag
- `prs`: PRS

Note: Columns `case`, `rv`, and
`prs` correspond to variables `Y`, `G`,
and `X` respectively in the causal pivot model.

##### 2.8.1 Pathogenic Variants

```
# mutate rare variant event counts into binary (>0) variable ####

events.df <- as.data.frame(as.matrix(gene.rv.events > 0))

# join cohort status, event status and prs data into one data frame ####

rv.prs.df <- bind_cols(
    cohort.flags,
    events.df[match(rownames(cohort.flags), rownames(events.df)),]
  ) %>%
  mutate(sample.id = rownames(.)) %>%
  inner_join(prs, by = 'sample.id') %>%
  select(sample.id, everything()) %>%
  as_tibble()

# create cohort dataframes list ####

cohort.dfs <- purrr::imap(COHORTS, function(cohort, cohort.name) {
  rv.prs.df[
    match(names(cohort$labels), rv.prs.df$sample.id),
    c('sample.id', cohort.name, cohort$gene.grp, cohort$prs)
  ] %>%
    magrittr::set_colnames(c('sample.id', 'case', 'rv', 'prs'))
})

# save results ####

saveRDS(rv.prs.df, file = file.path(OUT_DIR, 'rv.prs.df.rds'))
saveRDS(cohort.dfs, file = file.path(OUT_DIR, 'cohort.dfs.rds'))
```

##### 2.8.2 Control Variants

Re-create the primary analysis objects w/ the control variants.

```
# mutate rare variant event counts into binary (>0) variable ####

events.cv.df <- as.data.frame(as.matrix(gene.cv.events > 0))

# join cohort status, event status and prs data into one data frame ####

cv.prs.df <- bind_cols(
    cohort.flags,
    events.cv.df[match(rownames(cohort.flags), rownames(events.cv.df)),]
  ) %>%
  mutate(sample.id = rownames(.)) %>%
  inner_join(prs, by = 'sample.id') %>%
  select(sample.id, everything()) %>%
  as_tibble()

# create cohort dataframes list ####

cv.cohort.dfs <- purrr::imap(COHORTS, function(cohort, cohort.name) {
  cv.prs.df[
    match(names(cohort$labels), cv.prs.df$sample.id),
    c('sample.id', cohort.name, cohort$gene.grp, cohort$prs)
  ] %>%
    magrittr::set_colnames(c('sample.id', 'case', 'rv', 'prs'))
})

# save results ####

saveRDS(cv.prs.df, file = file.path(OUT_DIR, 'cv.prs.df.rds'))
saveRDS(cv.cohort.dfs, file = file.path(OUT_DIR, 'cv.cohort.dfs.rds'))
```

### 3 Rare Variant Summaries

#### 3.1 Create publishable variant Excel workbook

Export pathogenic rare variant data to Excel for supplemental
materials and analysis.

```
prs.cohort.summary.df <- purrr::imap(cohort.dfs, function(cohort.df, cohort.name) {
  gene.grp <- cohort.gene.grps[cohort.name]
  cohort.rv.marker.ids <- rv.flags[rv.flags$gene == gene.grp & rv.flags$pathogenic,]$marker.id
  cohort.gts <- as.data.frame(as.matrix(rv.mtx[cohort.df$sample.id,cohort.rv.marker.ids] ))
  n.control <- sum(!cohort.df$case)
  n.case <- sum(cohort.df$case)
  purrr::imap(cohort.gts, function(gts, marker.id) {
    cohort.df$gt <- gts[match(cohort.df$sample.id, rownames(cohort.gts))]
    
    freq.df <-  cohort.df %>%
      group_by(case) %>%
      summarize(f = mean(gt)) %>%
      ungroup()
    
    cohort.df %>%
      group_by(case) %>%
      filter(gt == 1) %>%
      summarize(prs.mean = mean(prs), n = n()) %>%
      ungroup() %>%
      mutate(marker.id = marker.id) %>%
      ungroup() %>%
      inner_join(freq.df, by = 'case')
  }) %>%
      do.call(bind_rows, .)
}) %>% concat.df.list(key.cols = 'cohort.name') %>%
  mutate(case = ifelse(case, 'case', 'control')) %>%
  pivot_wider(id_cols = c('cohort.name', 'marker.id'), names_from = 'case', values_from = c('prs.mean', 'f', 'n')) %>%
  select(
    `Marker` = 'marker.id',
    `Cohort` = 'cohort.name',
    `Mean PRS (Controls)` = 'prs.mean_control',
    `Mean PRS (Cases)` = 'prs.mean_case',
    `RV+ Count (Controls)` = 'n_control',
    `RV+ Count (Cases)` = 'n_case',
    `RV+ Frequency (Controls)` = 'f_control',
    `RV+ Frequency (Cases)` = 'f_case' 
  )

rv.summary.df <- annotations %>%
  filter(marker.id %in% rv.flags[rv.flags$pathogenic,]$marker.id) %>%
  mutate(cohort.af = apply(rv.mtx, 2, mean)[match(marker.id, colnames(rv.mtx))]) %>%
  select(
    `Marker` = marker.id,
    `Gene` = gene,
    `Transcript` = transcript,
    `Chrom` = chrom,
    `Position` = pos,
    `Ref` = ref,
    `Alt` = alt,
    `c.` = cchange,
    `a.` = achange,
    `gnomAD AF` = gnomad3.af_nfe,
    `UKB AF` = cohort.af,
    `ClinVar ID` = clinvar.allele_id,
    `Clinvar Significance` = clinvar.sig,
    `Clinvar Sign. Conflicting` = clinvar.sig_conf,
    `Revel` = revel.score,
    `CADD` = cadd_exome.phred,
  )

wb <- openxlsx::createWorkbook()
openxlsx::addWorksheet(wb, "Annotations")
openxlsx::writeData(wb, "Annotations", rv.summary.df)
openxlsx::addWorksheet(wb, "PRS + Counts")
openxlsx::writeData(wb, "PRS + Counts", prs.cohort.summary.df)
openxlsx::saveWorkbook(wb, file.path(OUT_DIR, 'rv.summary.xlsx'), overwrite = T)
```

#### 3.2 Plot variant classification summaries

##### 3.2.1 Variant mutation type waffle plot

Plot pathogenic rare variants by sequence ontology.

A waffle plot gives an intuitive feel for proportion of variants by
sequence ontology showing signals are not dominated by a single mutation
type (`MT`).

```
# create positive variants data frame w/ mutation types ####

mt.df <- left_join(
  filter(rv.flags, pathogenic),
  select(annotations, marker.id, mutation.type),
  by = 'marker.id'
) %>%
  mutate(mutation.type = as.factor(snakecase::to_title_case(mutation.type))) %>%
  group_by(gene, mutation.type) %>%
  summarize(n = n())  %>%
  ungroup() %>%
  mutate(gene = factor(gene, levels = gene.grps))

# create plot ####

# get sorted list of mutation type by most common - used to make color palette
mt.palette <- paletteer_d(
    "ggthemes::Classic_20",
    n = length(levels(mt.df$mutation.type))
  ) %>%
  purrr::set_names(levels(mt.df$mutation.type))

(
  mt.df %>%
    ggplot(aes(fill = mutation.type, values = n)) +
    geom_waffle(size = 0.33, colour = "white") +
    coord_equal() +
    facet_wrap(~gene) +
    theme_void() +
    theme(plot.margin = ggplot2::margin(2)) +
    labs(fill = 'Sequence Ontology')
) %>%
  save.plot('mutation_type.waffle', w = 7.35, h = 4.5 / 1.5, root.dir = PLOTS_DIR, sub.dir = 'supplementary')
```

```
saveRDS(mt.df, file.path(OUT_DIR, 'rv.mutation.type.rds'))
```

##### 3.2.2 Variant spread hisogram

Plot histogram summarizing variant mutation type and how variants are
distributed among cohorts. The x axis is the number of samples in which
a variant appears:

- bars to the left are rarer variants
- bars to the right are more common

The `max` annotation is the maximum number of samples in
which a single positive variant appears.

This plot further shows the causal pivot signal is not carried by a
single functional type.

```
# create dataframe counting positive samples per variants w/ SO annotation ####

alt.counts <- alt.sums.by.sample(rv.mtx, cohort.marks) %>%
  mtx.to.df('cohort', 'marker.id', 'alt.count') %>%
  left_join(select(annotations, marker.id, gene, mutation.type), by = 'marker.id') %>%
  mutate(mutation.type = as.factor(snakecase::to_title_case(mutation.type))) %>%
  rowwise() %>%
  filter(gene %in% COHORTS[[cohort]]$genes) %>%
  ungroup() %>%
  mutate(cohort = factor(cohort, levels = names(cohort.dfs)))

# calculate maximum number of count groups per cohort - used to place label
max.n <- alt.counts %>%
  group_by(cohort) %>%
  count(alt.count) %>%
  summarize(max.n = max(n)) %>%
  ungroup()

# create count group aggregation dataframe used for annotation
max.counts <- alt.counts %>%
  group_by(cohort) %>%
  summarize(max.count = max(alt.count)) %>%
  left_join(max.n, by = 'cohort') %>%
  rowwise() %>%
  mutate(label = paste("Max: ", max.count)) %>%
  ungroup()

# generate actual plot and save
mt.dist <- (
  alt.counts %>%
  mutate(n.samples = factor(ifelse(alt.count > 5, ">5", as.character(alt.count)), levels = c("1", "2", "3", "4", "5", ">5"))) %>%
  group_by(cohort, mutation.type, n.samples) %>%
  summarize(n.markers = n()) %>%
  ggplot(aes(x = n.samples, y = n.markers)) +
  geom_col(color = "black", aes(fill = mutation.type)) +
  scale_fill_manual(values = mt.palette) +
  facet_wrap(~cohort, scales='free_y') +
  theme_bw() +
  labs(
    x = 'Sample Counts',
    y = 'Num Markers',
    # title = paste("Variant Distribution Histogram w/ Sequence Ontology"),
    title = '',
    fill = "Mutation Type"
  ) +
  geom_text(data = max.counts, aes(label = label, y = max.n), x = 4)
) %>%
  save.plot('mutation_type.distribution', root.dir = PLOTS_DIR, sub.dir = 'supplementary')

mt.dist
```

```
# save results ####

# create useful rv count summary
rv.counts <- alt.counts %>%
  group_by(cohort, gene, alt.count) %>%
  summarize(n = n())

saveRDS(alt.counts, file = file.path(OUT_DIR, 'alt.counts.rds'))
saveRDS(rv.counts, file = file.path(OUT_DIR, 'rv.counts.rds'))
```

### 4 Perform MLE Permutation Testing

The Causal Pivot Maximum Likelihood Estimation (MLE) procedure is
applied to estimate gamma (the coefficient for G) and eta (the
coefficient for X \* G) of the reverse logistic model. A chi-squared test
statistic is calculated from the resulting maximum Log Likelihood Ratio
(LR). Empirical data and logistic regression is used to derive estimates
for model parameters.

MLE is performed via `optim()` using the build-in
`SANN` simulated annealing algorithm. In order to correct for
variability found in some areas of the parameter space, MLE is performed
multiple times and the median result is used.

Afterwards permutation analysis of the MLE statistic under G (the
response variable in the reverse model) is performed. The final test
statistic is a simple permutation z-statistic using the LR.

#### 4.1 Set Permutation Count

Lower permutation count reduces computation time, but increases
variability of results.

At least 256 permutations recommended for consistent results.

#### 4.2 Logit G

The equations for the Logit G model are given in equations.logitG.R.

```
# source MLE functions and load into local environment ####
source('../lrt_equations/equations.logitG.R')
list2env(lrt.logitG.equations, envir = environment())
```

```
## <environment: R_GlobalEnv>
```

```
source('utils/lrt.R')

n.mle.perms <- 1024 # number of permutations

# perform MLE testing under all cohorts ####

logitG.lrt.run <- purrr::map(cohort.dfs, function(cohort.df) {
  purrr::map(cohort.gene.grps, function(gene.grp) {
    # massage data
    G.df <- rv.prs.df[c('sample.id', gene.grp)] %>%
      magrittr::set_colnames(c('sample.id', 'G'))
    lrt.df <- cohort.df %>%
      left_join(G.df, by = 'sample.id') %>%
      mutate(
        Y = as.integer(case),
        G = as.integer(G),
        X = prs
      ) %>%
      select(Y, X, G)
  
    # calculate forward model parameter "targets"
    lrt.target.model <- broom::tidy(glm(Y ~ G * X, family = binomial(link = 'logit'), data = lrt.df))
    
    # estimate model parameters (alpha, beta, omega)
    lrt.est.model <- broom::tidy(glm(Y ~ X, family = binomial(link = 'logit'), data = lrt.df))
  
    params <- list(
      alpha = lrt.est.model$estimate[1],
      beta = lrt.est.model$estimate[2],
      gamma = 0,
      eta = 0,
      omega = mean(lrt.df$G) # rare variant frequency in disease
    )

    # calculate MLE w/ some jitter (internal to run.lrt) to avoid extreme results
    null.mle <- lapply(seq(16), function(i) {
      unlist(run.lrt(as.list(lrt.df), params, cases.only=T))
    }) %>% bind_rows()
    
    mle.run <- list(
      gamma = median(null.mle$gamma),
      eta = median(null.mle$eta),
      lr = median(null.mle$lr),
      p.val = median(null.mle$p.val)
    )
    
    fisher.information <- f.call(
      fisher.info,
      params %>%
        modifyList(mle.run) %>%
        modifyList(lrt.df)
    )
    
    # calculate permutation distribution of MLE results under G
    perm.result <- lapply(seq(n.mle.perms), function(i) {
      run.lrt(as.list(mutate(lrt.df, G = sample(G))), params, cases.only=T)
    }) %>%
      do.call(bind_rows, .) %>%
      as.data.frame()

    list(
      params = params,
      mle.target = lrt.target.model,
      mle.run = mle.run,
      fisher.information = fisher.information,
      perm.result = perm.result
    )
  }) %>%
    purrr::set_names(cohort.gene.grps)
})

c('params', 'mle.target', 'mle.run', 'perm.result') %>%
  purrr::set_names(paste('logitG', ., sep = '.')) %>%
  purrr::map(function(leaf.key) {
    concat.df.list(extract.leaf(logitG.lrt.run, leaf.key), key.cols = c('cohort.name', 'gene.grp')) %>%
      mutate(
        cohort.name = factor(cohort.name, levels = names(COHORTS)),
        gene.grp = factor(gene.grp, levels = cohort.gene.grps)
      )
  }) %>%
  list2env(mle.objs, envir = .GlobalEnv)
```

```
## <environment: R_GlobalEnv>
```

```
logitG.fisher.information <- extract.leaf(logitG.lrt.run, 'fisher.information')  

# sometimes the optimization produces unrealistic gamma/eta estimates
logitG.perm.result <- logitG.perm.result %>%
  group_by(cohort.name, gene.grp) %>%
  filter(
    eta > quantile(eta, 0.25) - 2 * IQR(eta),
    eta < quantile(eta, 0.75) + 2 * IQR(eta),
    gamma > quantile(gamma, 0.25) - 2 * IQR(gamma),
    gamma < quantile(gamma, 0.75) + 2 * IQR(gamma),
  ) %>%
  ungroup()

# calculate proportion of fisher information in cases
logitG.conf.int <- logitG.fisher.information %>%
  purrr::imap(function(a, cohort.name) {
    a %>%
      purrr::imap(function(b, gene.grp) {
        x <- solve(b$I1 * b$n1 + b$I2 * b$n2)
        y <- logitG.mle.run[
          logitG.mle.run$cohort.name == cohort.name &
            logitG.mle.run$gene.grp == gene.grp,]
        data.frame(
          gamma.low = y$gamma + qnorm(0.025, sd = sqrt(x[1,1])),
          gamma = y$gamma,
          gamma.high = y$gamma + qnorm(1 - 0.025, sd = sqrt(x[1,1])),
          eta.low = y$eta + qnorm(0.025, sd = sqrt(x[2,2])),
          eta = y$eta,
          eta.high = y$eta + qnorm(1 - 0.025, sd = sqrt(x[2,2]))
        )
      })
  }) %>%
  concat.df.list(key.cols = c('cohort.name', 'gene.grp'))

# calculate the 95th quantile of the LR permutation distribution
logitG.perm.q95 <- logitG.perm.result %>%
  group_by(cohort.name, gene.grp) %>%
  summarize(lr = q.95 <- quantile(lr, 0.95))

# calculate p.values for the cohorts based on 95th quantile of LR permutations
logitG.mle.p.vals <- inner_join(logitG.mle.run, logitG.perm.result, by = c('cohort.name', 'gene.grp'), suffix = c('.mle', '.perm')) %>%
  group_by(cohort.name, gene.grp) %>%
  summarize(p.val = mean(lr.perm > lr.mle))

# plot results ####

(
  logitG.perm.result %>%
  ggplot(aes(x = lr)) +
  facet_grid(cols = vars(cohort.name), rows = vars(gene.grp), scales = 'free') +
  geom_histogram(bins = 14) +
  geom_vline(aes(xintercept = lr), linewidth = 1, linetype = 'dashed', color = '#f42e3d', data = logitG.mle.run) +
  geom_vline(aes(xintercept = lr), linewidth = 1, linetype = 'dashed', data = logitG.perm.q95) +
  labs(
    y = 'Count',
    x = 'Log Likelihood',
  )
) %>%
  save.plot('logitG.MLE.permutation.histogram', root.dir = PLOTS_DIR, sub.dir = 'maximum.likelihood.estimation', w = 4.5 * 2, h = 4.5 * 2)
```

```
(
  mle.perm.logitG.contour.plt <- lapply(names(COHORTS), function(my.cohort.name) {
  c(
    lapply(gene.grps, function(my.gene.grp) {
      mle.perm <- logitG.perm.result %>%
        filter(gene.grp == my.gene.grp & cohort.name == my.cohort.name)
      mle.observed <- logitG.mle.run %>%
        filter(gene.grp == my.gene.grp & cohort.name == my.cohort.name)
      mle.target <- logitG.mle.target %>%
        filter(gene.grp == my.gene.grp & cohort.name == my.cohort.name) %>%
        filter(term %in% c('G', 'G:X')) %>%
        select(term, estimate) %>%
        pivot_wider(names_from = 'term', values_from = 'estimate') %>%
        select(gamma = G, eta = `G:X`)
      mle.perm %>%
        ggplot(aes(x = gamma, y = eta)) +
        geom_density_2d_filled(alpha = 1/3, bins = 7) +
        geom_density_2d(color = "black", alpha = 0.7, bins = 7) +
        # geom_point(shape = 1, size = 1/3, alpha = 1/3) +
        # geom_point(color = "black", size = 1, data = mle.observed) +
        geom_vline(xintercept = 0, linewidth = 0.5, linetype = 'dotted') +
        geom_hline(yintercept = 0, linewidth = 0.5, linetype = 'dotted') +
        geom_point(x = mle.target$gamma, y = mle.target$eta,
          color = "black", size = 4, shape = 4, stroke = 1) +
        geom_point(color = "#f42e3d", size = 3, shape = 16, data = mle.observed) +
        scale_fill_brewer(palette = 'PuBu') +
        labs(
          x = expression(gamma),
          y = expression(eta),
        ) +
        theme(legend.position = 'none')
    }),
    list(grid::textGrob(my.cohort.name, rot = -90))
  )
}) %>% 
  unlist(recursive = FALSE) %>%
  {
    c(map(gene.grps, ~grid::textGrob(.x)),
      list(plot_spacer()),
      .
    )
  } %>%
  wrap_plots(
    ncol = 4,
    byrow = T,
    heights = c(1, 5, 5, 5),
    widths = c(5, 5, 5, 1)
  ) +
  plot_layout(
    axis_titles = 'collect'
  )
) %>%
  save.plot('logitG.MLE.permutation.contour', root.dir = PLOTS_DIR, sub.dir = 'maximum.likelihood.estimation', w = 4.5 * 2, h = 4.5 * 2)
```

```
# save results ####
logitG.lrt.result <- list(
  params = logitG.params,
  mle.result = logitG.mle.run,
  mle.p.vals = logitG.mle.p.vals,
  perm.result = logitG.perm.result,
  fisher.information = logitG.fisher.information,
  logitG.conf.int = logitG.conf.int
)

saveRDS(logitG.lrt.result, file.path(OUT_DIR, 'logitG.mle.result.rds'))
```

#### 4.3 Liability G

The equations for the Logit G model are given in equations.liabilityG.R.

```
# source MLE functions and load into local environment ####
source('../lrt_equations/equations.liabilityG.R')
list2env(lrt.liabilityG.equations, envir = environment())
```

```
## <environment: R_GlobalEnv>
```

```
source('utils/lrt.R')

n.mle.perms <- 1024 # number of permutations

# perform MLE testing for HC190 ####

hc.cohort.df <- cohort.dfs$HC190 %>%
  left_join(select(cohort.meta, sample.id, ldl = ldl.inst.0), by = 'sample.id')

liabilityG.lrt.run <- purrr::map(unname(cohort.gene.grps), function(gene.grp) {
  # massage data
  G.df <- rv.prs.df[c('sample.id', gene.grp)]
  colnames(G.df) <- c('sample.id', 'G')
  lrt.df <- hc.cohort.df %>%
    left_join(G.df, by = 'sample.id') %>%
    mutate(
      y = (ldl - mean(ldl)) / sd(ldl),
      G = as.integer(G),
      X = prs
    ) %>%
    select(y, X, G)
  
  # calculate forward model paramter "targets"
  lrt.target.model <- broom::tidy(lm(y ~ G * X, data = lrt.df))

  # estimate model parameters (alpha, beta, delta, sige, omege)
  lrt.est.model <- lm(y ~ X, data = lrt.df)
  
  params <- list(
    alpha = tidy(lrt.est.model)$estimate[1],
    beta = tidy(lrt.est.model)$estimate[2],
    gamma = 0,
    eta = 0,
    delta = unname(quantile(lrt.df$y, prob = mean(hc.cohort.df < 4.9))),
    sige = sqrt(mean(lrt.est.model$residuals^2)),
    omega = mean(lrt.df$G)
  )
  
  # calculate MLE result
  mle.run <- run.lrt(as.list(lrt.df), params, cases.only=T)

  # calculate permutation distribution of MLE results under G
  perm.result <- lapply(seq(n.mle.perms), function(i) {
    r <- run.lrt(as.list(mutate(lrt.df, G = sample(G))), params, cases.only=T)
  }) %>%
    do.call(bind_rows, .) %>%
    as.data.frame()

  list(
    params = params,
    mle.target = lrt.target.model,
    mle.run = mle.run,
    perm.result = perm.result
  )
}) %>%
  purrr::set_names(cohort.gene.grps)

c('params', 'mle.target', 'mle.run', 'perm.result') %>%
  purrr::set_names(paste('liabilityG', ., sep = '.')) %>%
  purrr::map(function(leaf.key) {
    concat.df.list(extract.leaf(liabilityG.lrt.run, leaf.key), key.cols = c('gene.grp')) %>%
      mutate(gene.grp = factor(gene.grp, levels = cohort.gene.grps))
  }) %>%
  list2env(mle.objs, envir = .GlobalEnv)
```

```
## <environment: R_GlobalEnv>
```

```
# calculate the 95th quantile of the LR permutation distribution
liabilityG.perm.q95 <- liabilityG.perm.result %>%
  group_by(gene.grp) %>%
  summarize(lr = q.95 <- quantile(lr, 0.95, na.rm = T))

# calculate p.values for the cohorts based on 95th quantile of LR permutations
liabilityG.mle.p.vals <- inner_join(liabilityG.mle.run, liabilityG.perm.result, by = 'gene.grp', suffix = c('.mle', '.perm')) %>%
  group_by(gene.grp) %>%
  summarize(p.val = mean(lr.perm > lr.mle))

# plot results ####

(
  liabilityG.perm.result %>%
  ggplot(aes(x = lr)) +
  facet_wrap(~gene.grp, scales = 'free') +
  geom_histogram(bins = 14) +
  geom_vline(aes(xintercept = lr), linewidth = 1, linetype = 'dashed', color = '#f42e3d', data = liabilityG.mle.run) +
  geom_vline(aes(xintercept = lr), linewidth = 1, linetype = 'dashed', data = liabilityG.perm.q95) +
  labs(
    y = 'Count',
    x = 'Log Likelihood',
  )
) %>%
  save.plot('liabilityG.MLE.permutation.histogram', root.dir = PLOTS_DIR, sub.dir = 'maximum.likelihood.estimation')
```

```
(
  lapply(gene.grps, function(my.gene.grp) {
  mle.perm <- liabilityG.perm.result %>%
    filter(gene.grp == my.gene.grp)
  mle.observed <- liabilityG.mle.run %>%
    filter(gene.grp == my.gene.grp)
  mle.target <- liabilityG.mle.target %>%
    filter(gene.grp == my.gene.grp) %>%
    filter(term %in% c('G', 'G:X')) %>%
    select(term, estimate) %>%
    pivot_wider(names_from = 'term', values_from = 'estimate') %>%
    select(gamma = G, eta = `G:X`)
  mle.perm %>%
    ggplot(aes(x = gamma, y = eta)) +
    geom_density_2d_filled(alpha = 0.4, bins = 7) +
    geom_density_2d(color = "black", bins = 7) +
    # geom_point(shape = 1, size = 1/3, alpha = 1/3) +
    geom_vline(xintercept = 0, linewidth = 0.5, linetype = 'dotted') +
    geom_hline(yintercept = 0, linewidth = 0.5, linetype = 'dotted') +
    geom_point(x = mle.target$gamma, y = mle.target$eta,
      color = "black", size = 4, shape = 4, stroke = 1) +
    geom_point(color = "#f42e3d", size = 3, shape = 16, data = mle.observed) +
    scale_fill_brewer(palette = 'PuBu') +
    scale_x_continuous(limits = c(-1.5, 2)) +
    scale_y_continuous(limits = c(-1, 1)) +
    labs(
      x = expression(gamma),
      y = expression(eta),
    ) +
    theme(legend.position = 'none')
  }) %>%
    {
      c(., map(gene.grps, ~grid::textGrob(.x, rot = -90)) )
    } %>%
    wrap_plots(
      nrow = 3,
      byrow = F,
      widths = c(5, 1)
    ) +
    plot_layout(
      axis_titles = 'collect'
    )
) %>%
  save.plot('liabilityG.MLE.permutation.contour', root.dir = PLOTS_DIR, sub.dir = 'maximum.likelihood.estimation', w = 7.35 * 2 / 16 * 6, h = 4.5 * 2)
```

```
# save results ####
liabilityG.lrt.result <- list(
  params = liabilityG.params,
  mle.run = liabilityG.mle.run,
  mle.p.vals = liabilityG.mle.p.vals,
  perm.result = liabilityG.perm.result
  # fisher.information = liabilityG.fisher.information,
  # liabilityG.conf.int = liabilityG.conf.int
)

saveRDS(liabilityG.lrt.result, file.path(OUT_DIR, 'liabilityG.lrt.result.RDS'))
```

### 5 Plot Collider Effect

#### 5.1 Plot Case + Control Scatter Plot

Demonstrate unconditional (Case + Control / CC) independence of RV
status and PRS - a fundamental model assumption.

##### 5.1.1 Pathogenic Variants

```
# massage cohort.dfs into a single dataframe containing correctly paired RV/PRS ####

cc.prs.df <- concat.df.list(cohort.dfs, key.cols = 'cohort.name')
# get summary stats for plotting ####

cc.prs.stats.df <- cc.prs.df %>%
  group_by(cohort.name, rv) %>%
  summarize(
    mean.prs = mean(prs),
    err.prs = sqrt(var(prs)/n())
  ) %>%
  ungroup()

# calculate p-values using logistic regression ####

calc.p.val <- function(formula, data) { 
  summary(
    glm(formula,
      data = data,
      family = binomial(link='logit')
    )
  )[[12]][2,4]
}

cc.prs.pvals.df <- split(cc.prs.df, ~cohort.name) %>%
  map(function(prs.df) {
      tryCatch(calc.p.val(rv ~ prs, prs.df), error = default.na)
  }) %>%
  unlist() %>%
  tibble::enframe(name="cohort.name", value="p.val")

cc.prs.pvals.lbls <- cc.prs.pvals.df %>%
  rowwise() %>%
  mutate(stars = ifelse(p.val < 0.001, '***', ifelse(p.val < 0.01, '**', ifelse(p.val < 0.05, '*', 'n.s.'))))

cc.prs.pvals.df
```

```
## # A tibble: 3 × 2
##   cohort.name p.val
##   <chr>       <dbl>
## 1 BC          0.347
## 2 HC190       0.145
## 3 PD          0.641
```

```
# create plot ####

cc.prs.stats.df$cohort.name <- factor(cc.prs.stats.df$cohort.name, levels = names(cohort.dfs))

cc.scatter <- (
  cc.prs.stats.df %>%
  ggplot(aes(x = as.factor(rv), y = mean.prs, group = cohort.name)) +
  geom_point(position = position_dodge(width = 0.2), size = 2) +
  geom_errorbar(
    aes(ymin = mean.prs - err.prs, ymax = mean.prs + err.prs),
    position = position_dodge(width = 0.2),
    width = 0.1
  ) +
  geom_line(
    position = position_dodge(width = 0.2),
  ) +
  scale_x_discrete(labels = c("FALSE" = "RV-", "TRUE" = "RV+")) +
  scale_y_continuous(limits = c(-1, 1)) +
  facet_wrap(~cohort.name) +
  labs(
    x = "",
    y = "Mean PRS",
  ) +
  theme_bw() +
  theme(
    panel.grid = element_blank(),
    axis.text = element_text(size = 12, face = "bold"),
    axis.ticks = element_line(linewidth = 2),
    legend.position = "none",
    axis.title = element_text(size = 24, face = "bold"),
    strip.text = element_text(size = 11, face = "bold")
  ) +
  geom_text(data = cc.prs.pvals.lbls, aes(label = stars, y = 0.95), x = 2.25, vjust = 1, size = 6, show.legend=F)
) %>%
  save.plot('cc.scatter', w = 4.5 * 2, h = 3, root.dir = PLOTS_DIR, sub.dir = 'collider')

cc.scatter
```

##### Control Variants

Show the same for control variants.

```
# massage cohort.dfs into a single dataframe containing correctly paired RV/PRS ####

# control variant case+control
cv.cc.prs.df <- concat.df.list(cv.cohort.dfs, key.cols = 'cohort.name')

# get summary stats for plotting ####

cv.cc.prs.stats.df <- cv.cc.prs.df %>%
  group_by(cohort.name, rv) %>%
  summarize(
    mean.prs = mean(prs),
    err.prs = sqrt(var(prs)/n())
  ) %>%
  ungroup()

# calculate p-values using logistic regression ####

calc.p.val <- function(formula, data) { 
  summary(
    glm(formula,
      data = data,
      family = binomial(link='logit')
    )
  )[[12]][2,4]
}

cv.cc.prs.pvals.df <- split(cv.cc.prs.df, ~cohort.name) %>%
  map(function(prs.df) {
      tryCatch(calc.p.val(rv ~ prs, prs.df), error = default.na)
  }) %>%
  unlist() %>%
  tibble::enframe(name="cohort.name", value="p.val")

cv.cc.prs.pvals.lbls <- cv.cc.prs.pvals.df %>%
  rowwise() %>%
  mutate(stars = ifelse(p.val < 0.001, '***', ifelse(p.val < 0.01, '**', ifelse(p.val < 0.05, '*', 'n.s.'))))

cv.cc.prs.pvals.df 

# create plot ####

cv.cc.prs.stats.df$cohort.name <- factor(cv.cc.prs.stats.df$cohort.name, levels = names(cv.cohort.dfs))

cv.cc.prs.pvals.lbls$cohort.name <- factor(cv.cc.prs.pvals.lbls$cohort.name, levels = names(cv.cohort.dfs))

cv.cc.scatter <- (
  cv.cc.prs.stats.df %>%
  ggplot(aes(x = as.factor(rv), y = mean.prs, group = cohort.name)) +
  geom_point(position = position_dodge(width = 0.2), size = 2) +
  geom_errorbar(
    aes(ymin = mean.prs - err.prs, ymax = mean.prs + err.prs),
    position = position_dodge(width = 0.2),
    width = 0.1
  ) +
  geom_line(
    position = position_dodge(width = 0.2),
  ) +
  scale_x_discrete(labels = c("FALSE" = "RV-", "TRUE" = "RV+")) +
  scale_y_continuous(limits = c(-1, 1)) +
  facet_wrap(~cohort.name) +
  labs(
    x = "",
    y = "Mean PRS",
  ) +
  theme_bw() +
  theme(
    panel.grid = element_blank(),
    axis.text = element_text(size = 12, face = "bold"),
    axis.ticks = element_line(linewidth = 2),
    legend.position = "none",
    axis.title = element_text(size = 24, face = "bold"),
    strip.text = element_text(size = 11, face = "bold")
  ) +
  geom_text(data = cv.cc.prs.pvals.lbls, aes(label = stars, y = 0.95), x = 2.25, vjust = 1, size = 6, show.legend=F)
) %>%
  save.plot('cv.cc.scatter', w = 4.5 * 2, h = 3, root.dir = PLOTS_DIR, sub.dir = 'collider')

cv.cc.scatter
```

#### 5.2 Plot collider effect as PRS tertiles bar plot

Visualize the conditional causal collider effect in cases. Plot
proportion of samples w/ a RV event within each PRS tertile.

Plot is faceted on the three gene groups (columns) with correctly
paired cohort/PRS values. We expect statistically significant negative
association between RV and PRS along the diagonal (top-left to
bottom-right), where the cohort/PRS is paired with it’s causally similar
gene.

##### 5.2.1 Pathogenic Variants

```
# massage data w/ prs tertiles ####

n.events.df <- rv.prs.df %>%
  pivot_longer(cols = all_of(names(COHORTS)), names_to = 'cohort', values_to = 'in.cohort') %>%
  filter(in.cohort) %>%
  pivot_longer(cols = starts_with("PRS"), names_to = "prs.name", values_to = "value") %>%
  rowwise() %>%
  filter(prs.name == COHORTS[[cohort]]$prs) %>%
  ungroup() %>%
  pivot_longer(cols = all_of(names(gene.grps)), names_to = "gene.grp", values_to = "rv") %>%
  # anti_join(gene.excludes.df, by=c('sample.id', 'gene.grp')) %>%
  # unite(cohort, prs.name, col="cohort", sep='/') %>%
  select(sample.id, cohort, value, gene.grp, rv) %>%
  group_by(cohort, gene.grp) %>%
  mutate(prs.qnt = calc.quantile(value, n.qnt = 3, na.rm = T)) %>%
  ungroup() %>%
  filter(rv) %>%
  group_by(gene.grp, cohort, prs.qnt) %>% summarize(n = n())

n.events.df$gene.grp <- factor(n.events.df$gene.grp, levels=names(gene.grps))
n.events.df$cohort <- factor(n.events.df$cohort, levels=names(COHORTS))

# grab p.values from LRT

tertiles.p.vals <- expand.grid(cohort = names(COHORTS), gene.grp = unlist(gene.grps)) %>%
  rowwise() %>%
  mutate(p.val = logitG.lrt.run[[cohort]][[gene.grp]]$mle.run$p.val) %>%
  mutate(stars = ifelse(p.val < 0.001, '***', ifelse(p.val < 0.01, '**', ifelse(p.val < 0.05, '*', '')))) %>%
  left_join(
    n.events.df %>%
      group_by(cohort) %>% 
      summarize(y = max(n)
    ), by = c('cohort')
  )

# create and save plot ####
tertiles.3x3 <- (
  n.events.df %>%
  ggplot(aes(x = factor(prs.qnt), y = n)) +
  geom_bar(
    stat = "identity",
    color = 'black',
    fill = 'gray',
    width = 1
  ) +
  facet_grid(rows = vars(cohort), cols = vars(gene.grp), scales = "free_y") +
  scale_y_continuous(expand = expansion(mult = c(0, 0.5))) +
  geom_text(data = tertiles.p.vals, aes(label = stars, y = y), x = 3, vjust = -1/3, size = 6, show.legend=F) +
  theme_bw() +
  theme(
    panel.grid = element_blank(),
    axis.text = element_text(size = 12, face = "bold"),
    axis.ticks = element_line(linewidth = 1),
    legend.position = "none",
    axis.title = element_text(size = 24, face = "bold"),
    strip.text = element_text(size = 11, face = "bold")
  ) +
  labs(
    x = "PRS Tertile",
    y = "Num RV+ Samples",
  )
) %>%
   save.plot('3x3.tertiles', w = 4.5 * 2, h = 4.5 * 2, root.dir = PLOTS_DIR, sub.dir = 'collider')

tertiles.3x3
```

##### Control Variants

Plot the same for control variants.

```
# massage data w/ prs tertiles ####

n.cv.events.df <- cv.prs.df %>%
  pivot_longer(cols = all_of(names(COHORTS)), names_to = 'cohort', values_to = 'in.cohort') %>%
  filter(in.cohort) %>%
  pivot_longer(cols = starts_with("PRS"), names_to = "prs.name", values_to = "value") %>%
  rowwise() %>%
  filter(prs.name == COHORTS[[cohort]]$prs) %>%
  ungroup() %>%
  pivot_longer(cols = all_of(names(gene.grps)), names_to = "gene.grp", values_to = "rv") %>%
  # anti_join(gene.excludes.df, by=c('sample.id', 'gene.grp')) %>%
  # unite(cohort, prs.name, col="cohort", sep='/') %>%
  select(sample.id, cohort, value, gene.grp, rv) %>%
  group_by(cohort, gene.grp) %>%
  mutate(prs.qnt = calc.quantile(value, n.qnt = 3, na.rm = T)) %>%
  ungroup() %>%
  filter(rv) %>%
  group_by(gene.grp, cohort, prs.qnt) %>% summarize(n = n())

y <- expand.grid(cohort.name = names(cv.cohort.dfs), gene.grp = names(gene.grps))
x <- mapply(function(cohort.name, gene) {
  a <- cv.prs.df[,c(cohort.name, gene, COHORTS[[cohort.name]]$prs)] %>%
    magrittr::set_colnames(c('cohort', 'rv', 'prs')) %>%
    filter(cohort == T)
  p.val <- broom::tidy(glm(rv ~  prs, family = binomial(link = 'logit'), data = a))[2,5] %>% unlist()
  list(cohort.name = cohort.name, gene = gene, p.val = p.val)
}, paste(y$cohort.name), paste(y$gene.grp), SIMPLIFY=F) %>%
  do.call(bind_rows, .)

n.cv.events.df$gene.grp <- factor(n.cv.events.df$gene.grp, levels=names(gene.grps))
n.cv.events.df$cohort <- factor(n.cv.events.df$cohort, levels=names(COHORTS))

# create and save plot ####
cv.tertiles.3x3 <- (
  n.cv.events.df %>%
  ggplot(aes(x = factor(prs.qnt), y = n)) +
  geom_bar(
    stat = "identity",
    color = 'black',
    fill = 'gray',
    width = 1
  ) +
  facet_grid(rows = vars(cohort), cols = vars(gene.grp), scales = "free_y") +
  scale_y_continuous(expand = expansion(mult = c(0, 0.5))) +
  theme_bw() +
  theme(
    panel.grid = element_blank(),
    axis.text = element_text(size = 12, face = "bold"),
    axis.ticks = element_line(linewidth = 1),
    legend.position = "none",
    axis.title = element_text(size = 24, face = "bold"),
    strip.text = element_text(size = 11, face = "bold")
  ) +
  labs(
    x = "PRS Tertile",
    y = "Num RV+ Samples",
  )
) %>%
   save.plot('3x3.tertiles.controls', w = 4.5 * 2, h = 4.5 * 2, root.dir = PLOTS_DIR, sub.dir = 'collider')

cv.tertiles.3x3
```

### 6 Compose UKBB Collider Summary Plot

```
composed.theme <- theme(
  axis.title = element_text(face = 'bold', size = 10),
  axis.text = element_text(face = 'bold', size = 9),
  strip.text = element_text(face = 'bold', size = 12),
)

panel.a <- cc.scatter + composed.theme + scale_y_continuous(limits = c(-1, 1), breaks = c(-1, 0, 1))
panel.b <- tertiles.3x3 + composed.theme + scale_y_continuous(labels = function(x) x)
panel.c <- mt.dist + composed.theme +
  theme(
    legend.position = 'bottom',
    legend.box = 'horizontal',
    legend.text = element_text(face = 'bold', size = 12),
    legend.key.width = unit(0.25, 'cm')
  ) +
  guides(fill = guide_legend(nrow = 2))

(
  (((panel.a / panel.b) + plot_layout(heights = c(2, 8))) | panel.c) /
    guide_area() +
    plot_layout(heights = c(6, 1)) +
    plot_annotation(tag_levels = 'a', tag_suffix = ')') +
    plot_layout(guides = 'collect')
) %>%
  save.plot('fig3', root.dir = PLOTS_DIR, sub.dir = 'composed')
```

```
list(
  case.control.GvX = cc.prs.stats.df,
  tertile.pos.samples = n.events.df,
  rv.mt.alt.counts = alt.counts
) %>%
  saveRDS(file.path(OUT_DIR, 'fig3.RDS'))
```

### 7 Perform Ancestry Confounder Control Analysis

#### 7.1 Ancestry k Nearest Neighbors analysis

The causal collider effect is shown when controlling for
ancestry.

Ancestry is controlled for by observing the causal collider effect
between all cases and the (k=2) closest controls in the ancestry PCA
space.

A permutation z-statistic is calculated under G (the response
variable in the reverse model).

A tSNE projection of the PCA space is used to visualize the
procedure.

Note: in tSNE space, observations may appear more distant than they
really are and are shown purely as an intuitive aid.

```
# set parameters ####

n = 1000 # the number of permutations when creating permutation distributions
k = 2 # number of RV- neighborsto use when calculating PRS delta
r = 0.637 # the "decay" factor to simulate decreasing eigenvalues of PCs

tsne.seed = 5 # RNG seed for tSNE
x.lim <- c(-15, 15) # x limits for tSNE zoom plot
y.lim <- c(-15, 15) # y limits for tSNE zoom plot

# extract first 5 PCs to calculate nearest neighbors ####

pc.cols <- paste0('pc.', seq(5))
knn.pcs <- cohort.meta %>%
    select(sample.id, all_of(pc.cols))

# iterate over cohorts and perform analysis #####

knn.results <- cohort.dfs %>%
  purrr::imap(function(cohort.df, cohort.name) {
    # helper function to find nearest neighbors ####
    pos.neg.knn <- function(pc.distances, k, rv) {
      pos.idx <- which(rv)
      neg.idx <- which(!rv)
      
      distances::nearest_neighbor_search(
        distances = pc.distances,
        k = k,
        query_indices = pos.idx,
        search_indices = neg.idx
      )
    }
    
    # helper function to calculate test statistic ####
    knn.collider.prs.diff.mean <- function(knn) {
      prs.diffs <- sapply(seq(ncol(knn)), function(idx) {
        q.idx <- as.integer(colnames(knn)[idx])
        h.idx <- knn[,idx]
        rv.pcs.df[q.idx,]$prs - mean(rv.pcs.df[h.idx,]$prs)
      })
      
      mean(prs.diffs)
    }
  
    # rv/prs dataframe w/ PCs ####
    rv.pcs.df <- cohort.df %>%
      filter(case) %>%
      select(sample.id, rv, prs) %>%
      inner_join(knn.pcs, by = 'sample.id')
    
    # calculate distances in PC space ####
    pcs.mat <- as.matrix(select(rv.pcs.df, starts_with('pc')))
    rownames(pcs.mat) <- rv.pcs.df$sample.id
    decay <- exp(-r * 0:(ncol(pcs.mat) - 1))
    weights <- decay / sum(decay)
    pc.distances <- distances::distances(pcs.mat, weights=weights)
    
    # calculate the observed statistic ####
    prs.diff <- knn.collider.prs.diff.mean(pos.neg.knn(pc.distances, k, rv.pcs.df$rv))
    
    # create permutation distribution of test statistic ####
    prs.diff.sdsm <- sapply(seq(n), function(x) {
      knn.collider.prs.diff.mean(pos.neg.knn(pc.distances, k, sample(rv.pcs.df$rv)))
    })
    
    # store permutation results ####
    perm.res <- list(
      mean.diff = prs.diff,
      perm.sdsm = prs.diff.sdsm
    )
    
    # calculate 5 nearest neighbors and create tSNE (for vizualiation) ####
    set.seed(tsne.seed)
    knn <- pos.neg.knn(pc.distances, k = 5, rv.pcs.df$rv)
    tsne <- Rtsne(as.matrix(as.dist(pc.distances)), is_distance = T)
    
    list(
      rv.pcs.df = rv.pcs.df,
      pc.distances = pc.distances,
      knn = knn,
      tsne = tsne,
      perm.res = perm.res
    )
})
  
knn.plots <- knn.results %>%
  purrr::map(function(results) {
    # helper function to create tSNE plots
    make.tsne.plots <- function(tsne, knn, rv.pcs.df, x.lim = c(-30, 30), y.lim = c(-30, 30), min.alpha = 0.25) {
      tsne.df <- cbind(
        rv.pcs.df[,c('sample.id', 'rv')],
        data.frame(tSNE.1 = tsne$Y[,1], tSNE.2 = tsne$Y[,2])
      ) %>%
        mutate(rv = factor(ifelse(rv, 'RV+', 'RV-'), levels = c('RV-', 'RV+')))
      
      knn.connections.df <- do.call(
        rbind,
        apply(knn, 1, function(row) {
          x.idx <- as.integer(colnames(knn))
          y.idx <- row
          data.frame(
            from = tsne.df$sample.id[x.idx],
            to = tsne.df$sample.id[y.idx],
            from.x = tsne.df$tSNE.1[x.idx],
            to.x = tsne.df$tSNE.1[y.idx],
            from.y = tsne.df$tSNE.2[x.idx],
            to.y = tsne.df$tSNE.2[y.idx]
          )
        })
      ) %>%
        filter(
          from.x > x.lim[1] & from.x < x.lim[2] & to.x > x.lim[1] & to.x < x.lim[2]
          & from.y > y.lim[1] & from.y < y.lim[2] & to.y > y.lim[1] & to.y < y.lim[2]
        )
      
      full.plot <- tsne.df %>%
        arrange(rv) %>%
        ggplot(aes(x = tSNE.1, y = tSNE.2, color = rv, alpha = rv, size = rv)) +
        geom_point() +
        scale_color_manual(values = c('RV+' = "#f42e3d", 'RV-' = "black")) +
        scale_size_manual(values = c('RV+' = 4.5, 'RV-' = 1.5)) +
        scale_alpha_manual(values = c('RV+' = 1.0, 'RV-' = 1)) +
        labs(x = "tSNE 1", y = "tSNE 2") +
        theme_classic() +
        theme(
          axis.text = element_blank(),
          axis.ticks = element_blank(),
          legend.position = "none",
          axis.title = element_text(size = 30, face = "bold")
        )
      
      knn.plot <- tsne.df %>%
        ggplot() +
        geom_segment(
          data = knn.connections.df,
          aes(x = from.x, y = from.y, xend = to.x, yend = to.y),
          linetype = 'dashed',
          linewidth = 0.5
        ) +
        geom_point(
          data = tsne.df %>%
            filter(sample.id %in% c(knn.connections.df$from, knn.connections.df$to)) %>%
            filter(`tSNE.1` > x.lim[1] & `tSNE.1` < x.lim[2] & `tSNE.2` > y.lim[1] & `tSNE.2` < y.lim[2]) %>%
            arrange(rv),
          aes(x = tSNE.1, y = tSNE.2, color = rv),
          size = 8
        ) +
        scale_color_manual(values = c('RV+' = "#f42e3d", 'RV-' = "black")) +
        xlim(x.lim) +
        ylim(y.lim) +
        labs(x = "", y = "") +
        theme_void() +
        theme(
          legend.position = "none"
        )
        
        list(full.plot, knn.plot)
    }
    
    # helper function to create histogram plot
    plot.hist.w.vline <- function(pc.distances, mean.diff, perm.sdsm, q.05) {
      data.frame(x = perm.sdsm) %>%
        ggplot(aes(x = x)) +
        geom_histogram(
          aes(y = ..density.., fill = x < q.05),
          bins = 15,
          fill = 'gray',
          color = 'black',
          alpha = 0.7,
          position = "identity"
        ) +
        scale_x_continuous(expand = c(0,0)) +
        scale_y_continuous(expand = c(0,0)) +
        geom_vline(xintercept = mean.diff, linetype = "dashed", color = "#f42e3d", linewidth = 3) +
        geom_vline(xintercept = q.05, linetype = "dashed", color = "black", linewidth = 1.5) +
        labs(
          x = "Mean PRS Diff",
          y = ""
        ) +
        theme_classic() +
        theme(
          axis.line.y = element_blank(),
          axis.text.y = element_blank(),
          axis.ticks.y = element_blank(),
          axis.text.x = element_text(size = 20, face = "bold"),
          axis.ticks.x = element_line(linewidth = 2),
          legend.position = "none",
          axis.title = element_text(size = 30, face = "bold")
        )
    }
    
    # extract objects from results
    rv.pcs.df <- results$rv.pcs.df
    pc.distances <- results$pc.distances
    knn <- results$knn
    tsne <- results$tsne
    mean.diff <- results$perm.res$mean.diff
    perm.sdsm <- results$perm.res$perm.sdsm
    
    # create tSNE plots
    tsne.plots <- make.tsne.plots(tsne, knn, rv.pcs.df, x.lim = x.lim, y.lim = y.lim)
    
    # perform permutation t.test
    sdsm.mean <- mean(perm.sdsm)
    sdsm.sd <- sd(perm.sdsm)
    
    z.score <- (mean.diff - sdsm.mean) / sdsm.sd
    p.val <- pnorm(z.score)
  
    q.05 <- quantile(perm.sdsm, 0.05)
    
    hist.plot <- plot.hist.w.vline(pc.distances, mean.diff, perm.sdsm, q.05)
    
    list(
      full.tsne = tsne.plots[[1]],
      zoom.tsne = tsne.plots[[2]],
      hist = hist.plot,
      results = list(
        sdsm.mean = sdsm.mean,
        sdsm.sd = sdsm.sd,
        mean.diff = mean.diff,
        q.05 = q.05,
        z.score = z.score,
        p.val = p.val
      )
    )
})

# save plots ####

purrr::imap(knn.plots, function(plots, cohort.name) {
  save.plot(plots$full.tsne, paste(cohort.name, 'full.tsne', sep='.'), w = 4.5 * 2, h = 4.5 * 2, root.dir = PLOTS_DIR, sub.dir = 'ancestry.balancing/k.nearest.neighbors')
  save.plot(plots$zoom.tsne, paste(cohort.name, 'zoom.tsne.knn', sep='.'), w = 4.5 * 2, h = 4.5 * 2, root.dir = PLOTS_DIR, sub.dir = 'ancestry.balancing/k.nearest.neighbors')
  save.plot(plots$hist, paste(cohort.name, 'knn.hist.t', sep='.'), w = 4.5 * 2, h = 4.5 * 2, root.dir = PLOTS_DIR, sub.dir = 'ancestry.balancing/k.nearest.neighbors')
})
```

```
## $HC190
```

```
## 
## $BC
```

```
## 
## $PD
```

```
knn.plots %>% map(function(i) i$results$p.val)
```

```
## $HC190
## [1] 5.072583e-08
## 
## $BC
## [1] 0.004241831
## 
## $PD
## [1] 0.008170327
```

```
# save results ####

saveRDS(modifyList(knn.results, knn.plots), file = file.path(OUT_DIR, 'fig4.rds'))

# display plots ####

knn.plots$HC190$full.tsne
```

```
knn.plots$HC190$zoom.tsne
```

```
knn.plots$HC190$hist
```

```
knn.plots$BC$full.tsne
```

```
knn.plots$BC$zoom.tsne
```

```
knn.plots$BC$hist
```

```
knn.plots$PD$full.tsne
```

```
knn.plots$PD$zoom.tsne
```

```
knn.plots$PD$hist
```

#### 7.2 Ancestry Balancing via Inverse Probability Weighting

A lateral method for ancestry balancing is performed using inverse
probability weighting. Propensities are calculated using random forests
over the first 15 ancestry PCs from UKB. Again a permutation z-statistic
is calculated under G; the statistic is the difference in weighted PRS
sums between RV+ and RV- cases.

Random forests “importance” metrics are plotted as supplemental
information.

```
# set parameters ####

n.trees <- 1000 # number of trees in forest
n.perms <- 10000 # number of permutations for permutation testing
n.pcs <- 15 # number of principal components to use as features

# extract ancestry PCs into dataframe - for simplicity ####

pcs.df <- cohort.meta %>%
  select(sample.id, starts_with('pc.'))

# create vector of PCs we'll be extracting to train model ####
# note: the order is important due to our decay function

pc.cols <- paste0('pc.', seq(n.pcs))

# ancestry - inverse probability weighting results
a.ipw.results <- purrr::map(cohort.dfs, function(cohort.df) {
  # join cohort data to ancestry principal components (there are 40 in total)
  x <- inner_join(
    cohort.df,
    pcs.df,
    by = 'sample.id'
  ) %>%
    filter(case)
  
  # train model w/ importance
  my.rfm <- randomForest(
    x = x[,pc.cols,drop=F],
    y = as.factor(x$rv),
    ntree = n.trees,
    importance = T,
  )
  
  # store predictor importance metrics
  imp <- as.data.frame(importance(my.rfm))
  
  # set propensity score from model prediction
  x$prop.score <- predict(my.rfm, x[,pc.cols,drop=F], type = "prob")[,'TRUE']
  
  # calculate permutation distribution by permuting PRS and re-weighting
  p.dist <- sapply(1:n.perms, function(.) {
    # permute prs and weight by propensity score (or 1 - score if event == F)
    prs.sampled <- sample(x$prs) / ifelse(x$rv == T, x$prop.score, (1 - x$prop.score))
    # calculate test statistic - the normalized difference of weighted PRS b/w RV+ and RV- samples
    (sum(prs.sampled[which(x$rv==T)]) - sum(prs.sampled[which(x$rv==F)])) / nrow(x)
    # mean(prs.sampled[which(x$event==T)]) - mean(prs.sampled[which(x$event==F)])
  })
  
  # calculate observed test statistic
  x$prs.weighted <- x$prs / ifelse(x$rv == T, x$prop.score, (1 - x$prop.score))
  t <- (sum(x[x$rv==T,]$prs.weighted) - sum(x[x$rv==F,]$prs.weighted)) / nrow(x)
  
  u <- mean(p.dist) # mean of permutation distribution
  o <- sd(p.dist) # standard deviation of permutation distribution
  z <- (t - u) / o # z score of observed statistic
  p.val <- pnorm(z) # p.value of test
  
  p.dist.std <- (p.dist - u) / o # standardized distribution
  
  list(
    p.dist.std = p.dist.std,
    z = z,
    imp = imp,
    p.val = p.val
  )
})

# create plots
for (cohort.name in names(COHORTS)) {
  r <- a.ipw.results[[cohort.name]]
  
  p.dist.std <- r$p.dist.std
  z <- r$z
  imp <- r$imp
  
  # plot permutation test histogram w/ line at observed statistic
  hist.plt <- data.frame(x = p.dist.std) %>%
    ggplot(aes(x = x)) +
    geom_histogram(
      aes(y = ..density..),
      bins = 15,
      fill = 'gray',
      color = 'black',
      alpha = 0.7,
      position = "identity"
    ) +
    geom_vline(xintercept = z, linetype = "dashed", color = "red", size = 1.5) +
    geom_vline(xintercept = quantile(p.dist.std, 0.05), linetype = "dashed", color = "black", size = 1.5) +
    scale_x_continuous(expand = c(0,0)) +
    scale_y_continuous(expand = c(0,0)) +
    theme_classic() +
    theme(
      axis.line.y = element_blank(),
      axis.text.y = element_blank(),
      axis.ticks.y = element_blank(),
      axis.text.x = element_text(size = 12, face = "bold"),
      axis.ticks.x = element_line(linewidth = 2),
      legend.position = "none",
      axis.title = element_text(size = 30, face = "bold")
    ) +
    labs(
      x = "",
      y = ""
    )
  
  # define helper function to create predictor importance plots
  plot_imp <- function(imp, measure) {
    # importance(my.rfm) returns multiple importance measure
    # we are primarily concerned with MeanDecreaseAccuracy and MeanDecreaseGini
    # they quantify the average decrease in two different measures of predictor
    # performance when predictors are REMOVED - so higher decrease is better
    imp$predictor <- rownames(imp)
    
    imp$measure <- imp[[measure]]
    # we make predictor factor with levels sorted to make ggplot2 plot them in correct order
    imp$predictor <- factor(imp$predictor, levels = imp$predictor[order(imp$measure)])
    
    imp %>%
      select(predictor, measure) %>%
      arrange(desc(predictor)) %>%
      ggplot(aes(y = predictor, x = measure)) +
      geom_hline(aes(yintercept=predictor), linetype = 'dotted', linewidth = 0.5, alpha = 0.33) +
      geom_point(shape = 21, size = 5, fill = 'white') +
      theme_classic() +
      theme(
        axis.ticks.y = element_blank(),
        axis.line.x = element_line(),
        legend.position = "none",
      )
  }
  
  rownames(imp) <- gsub('pc.', 'PC', rownames(imp))
  # make importance accuracy plot
  imp.acc.plt <- plot_imp(imp, 'MeanDecreaseAccuracy') +
      labs(
        y = "",
        x = ""
      )
  
  # make importance gini purity plot
  imp.gini.plt <- plot_imp(imp, 'MeanDecreaseGini') +
      labs(
        y = "",
        x = ""
      )
  
  save.plot(r$hist.plt, paste(cohort.name, 'perm.hist', sep='.'), w = 4.5 * 2, h = 4.5 * 2, root.dir = PLOTS_DIR, sub.dir = 'ancestry.balancing/inverse.probability.weighting')
  save.plot(r$imp.acc.plt, paste(cohort.name, 'imp.acc', sep='.'), w = 4.5, h = 4.5 * 2, root.dir = PLOTS_DIR, sub.dir = 'ancestry.balancing/inverse.probability.weighting')
  save.plot(r$imp.gini.plt, paste(cohort.name, 'imp.gini', sep='.'), w = 4.5, h = 4.5 * 2, root.dir = PLOTS_DIR, sub.dir = 'ancestry.balancing/inverse.probability.weighting')
  
  # store results
  r$hist.plt <- hist.plt
  r$imp.acc.plt <- imp.acc.plt
  r$imp.gini.plt <- imp.gini.plt
  
  a.ipw.results[[cohort.name]] <- r
}
```

```
## NULL
## NULL
## NULL
## NULL
## NULL
## NULL
## NULL
## NULL
## NULL
```

```
# composed plot

composed.theme <- theme(
  axis.title.y = element_text(size = 12, face = "bold"),
  axis.title.x = element_text(size = 11, face = "bold"),
  axis.text.y = element_text(size = 12, face = "bold"),
  axis.text.x = element_text(size = 12, face = "bold"),
  axis.ticks.x = element_line(linewidth = 2),
)

# row 1 (HC190)

a.ipw.plts <- lapply(names(COHORTS), function(cohort.name) {
  r <- a.ipw.results[[cohort.name]]
  list(
    grid::textGrob(cohort.name, rot = 90, gp = grid::gpar(fontface = 'bold')),
    r$hist.plt,
    r$imp.acc.plt,
    r$imp.gini.plt
  )
})%>%
  unlist(recursive = F)

a.ipw.plts[[10]] <- a.ipw.plts[[10]] + labs(x = "Weighted PRS Delta")
a.ipw.plts[[11]] <- a.ipw.plts[[11]] + labs(x = "Importance (Accuracy)")
a.ipw.plts[[12]] <- a.ipw.plts[[12]] + labs(x = "Importance (Gini)")

a.ipw.composed.plt <- a.ipw.plts %>%
  wrap_plots(
    nrow = 3,
    byrow = T,
    widths = c(1, 6, 4, 4)
  ) %>%
  {. & composed.theme}

a.ipw.composed.plt
```

```
save.plot(a.ipw.composed.plt, paste('ancestry.ipw', sep='.'), w = 6 * 2, h = 4.5 * 2, root.dir = PLOTS_DIR, sub.dir = 'ancestry.balancing/inverse.probability.weighting')
```

```
purrr::map(a.ipw.results, ~.$p.val)
```

```
## $HC190
## [1] 2.131726e-11
## 
## $BC
## [1] 0.0007561405
## 
## $PD
## [1] 0.02214512
```

```
saveRDS(a.ipw.results, file = file.path(OUT_DIR, 'a.ipw.results.rds'))
```

### 8 Supplementary LSD/PD “Genetic Load” Analysis

Show the causal collider signal is still present when considering RV
status as an ordinal variable (as opposed to a binary variable). We
consider the “genetic load” of rare deleterious and ClinVar pathogenic
variants within a set of genes related to Lysosomal Storage Disease. We
then observe the collider signal between this variable and PD PRS in PD
cases.

TODO: reasons for doubletons

```
# classify pathogenic variants in the LSD gene list ####

lsd.rv.flags <- annotations %>%
  filter(gene %in% lsd.genes) %>%
  mutate(
    class.rare = gnomad3.af_nfe < gnomad.cutoff,
  ) %>%
  classify.clinvar() %>%
  classify.lof() %>%
  class.na.false() %>%
  mutate(
    class.positive = class.clinvar.pathogenic | (class.rare & class.loss.of.function)
  ) %>%
  select(marker.id, gene, class.positive)

# remove variants previously associated w/ PRS ####

lsd.rv.flags$pathogenic <- lsd.rv.flags$class.positive &
  !(lsd.rv.flags$marker.id %in% prs.sig.marker.ids)

# get positive LSD gene named vector for GT aggregation ####

lsd.pos <- lsd.rv.flags %>%
  filter(pathogenic & gene %in% lsd.genes) %>%
  with(setNames(gene, marker.id))

# calculate RV status for each LSD gene ####

lsd.events <- alt.sums.by.var(
  geno.mtx[,match(names(lsd.pos), colnames(geno.mtx))] > 0,
  lsd.pos
)

# calculate total number of LSD events per sample ####

lsd.df <- cbind(cohort.dfs$PD, n = rowSums(lsd.events)[match(cohort.dfs$PD$sample.id, rownames(lsd.events))])

# create PD/LSD genetic load collider boxplot ####

lsd.boxplot <- lsd.df %>%
  filter(case) %>%
  mutate(n = as.factor(map_chr(n, ~ ifelse(.x < 2, as.character(.x), '2+')))) %>%
  ggplot(aes(x = n, y = prs)) +
  geom_boxplot(fill = 'gray') +
  theme_bw() +
  theme(
    axis.text.x = element_text(size = 14, face = "bold"),
    axis.text.y = element_text(size = 14, face = "bold"),
    axis.title = element_text(size = 17, face = "bold"),
  ) +
  labs(
    y = "PRS",
    x = "Pathogenic Burden"
  )


save.plot(lsd.boxplot, 'boxplot', w = 4.5 * 2, h = 4.5 * 2,  root.dir = PLOTS_DIR, sub.dir = 'LSD.burden')
```

```
# logistic model of genetic load collider signal ####

lsd.burden.model <- tidy(glm(n ~ prs, family = poisson(link = "log"), data = lsd.df[lsd.df$case,]))

lsd.burden.model
```

```
## # A tibble: 2 × 5
##   term        estimate std.error statistic   p.value
##   <chr>          <dbl>     <dbl>     <dbl>     <dbl>
## 1 (Intercept)   -2.30     0.0615    -37.4  2.43e-305
## 2 prs           -0.147    0.0597     -2.47 1.35e-  2
```

```
# analyze doubletons ####

pd.sample.ids <- cohort.dfs$PD[cohort.dfs$PD$case,]$sample.id

# calculate pairwise matrix of num samples w/ genetic load == 2 (doubletons) ####

lsd.dbl <-  lsd.events[rowSums(lsd.events) == 2,]
lsd.dbl <- lsd.dbl[na.omit(match(pd.sample.ids, rownames(lsd.dbl))),]
lsd.dbl <- lsd.dbl[,colSums(lsd.dbl) > 0]
lsd.pairs <- lsd.dbl %>% apply(2, function(g1) {
  apply(lsd.dbl, 2, function(g2) {
    sum(g1 & g2)
  })
})

lsd.pairs.long <- as.data.frame(lsd.pairs) %>%
  mutate(g1 = rownames(.)) %>%
  pivot_longer(cols = -g1, names_to = 'g2', values_to = 'n')

# create doubleton heatmap ####

lsd.dbl.heatmap <- lsd.pairs.long %>%
  ggplot(aes(x = g1, y = g2, fill = n)) +
  geom_tile(aes(width = 1, height = 1), fill = NA, color = "grey50") +
  geom_tile(width = 0.8, height = 0.8) +
  scale_fill_gradientn(
    colours = c("white", "#f42e3d"),
    breaks = 0:max(lsd.pairs.long$n)
  ) +
  theme_bw() +
  theme(
    aspect.ratio = 1,
    panel.grid = element_blank(),
    axis.text.x = element_text(size = 12, face = "bold", angle = 45, hjust = 1),
    axis.text.y = element_text(size = 12, face = "bold"),
    axis.title = element_text(size = 17, face = "bold"),
  ) +
  labs(
    x = 'Gene 1',
    y = 'Gene 2',
  )

save.plot(lsd.dbl.heatmap, 'heatmap', w = 4.5 * 2, h = 4.5 * 2, root.dir = PLOTS_DIR, sub.dir = 'LSD.burden')
```

```
(
  (lsd.boxplot + lsd.dbl.heatmap) +
    plot_annotation(tag_levels = 'a', tag_suffix = ')')
) %>%
  save.plot('fig5', root.dir = PLOTS_DIR, sub.dir = 'composed')
```

```
# save results ####

list(
  df = lsd.df,
  lsd.glm = lsd.burden.model,
  lsd.pairs = lsd.pairs
) %>%
  saveRDS( file = file.path(OUT_DIR, 'fig5.rds'))
```
